# Potent antitumor activity of a designed interleukin-21 mimic

**DOI:** 10.1101/2024.12.06.626481

**Authors:** Jung-Ho Chun, Birkley S. Lim, Suyasha Roy, Michael J. Walsh, Gita C. Abhiraman, Kevin Zhangxu, Tavus Atajanova, Or-Yam Revach, Elisa C. Clark, Peng Li, Claire A. Palin, Asheema Khanna, Samantha Tower, Rakeeb Kureshi, Megan T. Hoffman, Tatyana Sharova, Aleigha Lawless, Sonia Cohen, Genevieve M. Boland, Tina Nguyen, Frank Peprah, Julissa G. Tello, Samantha Y. Liu, Chan Johng Kim, Hojeong Shin, Alfredo Quijano-Rubio, Kevin M. Jude, Stacey Gerben, Analisa Murray, Piper Heine, Michelle DeWitt, Umut Y. Ulge, Lauren Carter, Neil P. King, Daniel-Adriano Silva, Hao Yuan Kueh, Vandana Kalia, Surojit Sarkar, Russell W. Jenkins, K. Christopher Garcia, Warren J. Leonard, Michael Dougan, Stephanie K. Dougan, David Baker

**Author notes:** Corresponding authors. Mail to, or. These authors contributed equally to this work (Jung-Ho Chun, Birkley S. Lim, Suyasha Roy, Michael J. Walsh, Gita C. Abhiraman).

## Abstract

Long-standing goals of cancer immunotherapy are to activate cytotoxic antitumor T cells across a broad range of affinities while dampening suppressive regulatory T (Treg) cell responses, but current approaches achieve these goals with limited success. Here, we report a *de novo* IL-21 mimic, 21h10, designed to have augmented stability and high signaling potency in both humans and mice. In multiple animal models and in *ex vivo* human melanoma patient derived organotypic tumor spheroids (PDOTS), 21h10 showed robust antitumor activity. 21h10 generates significantly prolonged STAT signaling *in vivo* compared with native IL-21, and has considerably stronger anti-tumor activity. Toxicities associated with systemic administration of 21h10 could be mitigated by TNFα blockade without compromising antitumor efficacy. In the tumor microenvironment, 21h10 induced highly cytotoxic antitumor T cells from clonotypes with a range of affinities for endogenous tumor antigens, robustly expanding low-affinity cytotoxic T cells and driving high expression of interferon-𝛾 (IFN-𝛾) and granzyme B compared to native IL-21, while increasing the frequency of IFN-𝛾^+^ Th1 cells and reducing that of Foxp3^+^ Tregs. As 21h10 has full human/mouse cross-reactivity, high stability and potency, and potentiates low-affinity antitumor responses, it has considerable translational potential.

## Main

IL-21 plays a key role in the differentiation of effector T cells^1–4^, T follicular helper cells (T_FH_)^5^, B cells^6,7^, natural killer (NK) cells^3^, macrophages^8^, and dendritic cells^9,10^. IL-21 promotes the proliferation and survival of CD8^+^ T cells and the generation of memory CD8^+^ T cells^10–13^. In addition, IL-21 enhances the effector functions of CD8^+^ T cells^10,14^ and cooperates with IL-15 to further augment the expansion of these cells^4^. Multiple murine tumor models and early-stage clinical trials^15,16^ of recombinant IL-21 have been completed in several types of cancer, including melanoma^17–22^, renal cell carcinoma^21–23^, ovarian cancer^24^, and non-Hodgkin’s lymphoma^25^, both as a single agent and in combination with checkpoint inhibitors and cancer vaccines. IL-21 was also shown to be more effective than IL-2 when PMEL-1 TCR transgenic CD8^+^ T cells were cultured *in vitro* with cytokines prior to adoptive transfer into B16F10 melanoma-bearing mice^26^. Although the initial clinical testing of IL-21-based cancer immunotherapies has been encouraging, the precise effect of IL-21 on antitumor responses remains incompletely understood. Despite its promise, IL-21 has low stability, resulting in suboptimal pharmacokinetic properties, and the limited antitumor activity of human IL-21 (hIL-21) in mice has complicated the use of animal models to predict the toxicity and activity of candidate IL-21 therapeutics in humans^27–31^.

Inspired by previous success in designing *de novo* IL-2 mimics for cancer immunotherapy^32,33^, we reasoned that a *de novo* protein mimic of IL-21 with augmented stability that potently induces signaling in both humans and mice could overcome the limitations of native IL-21. We set out to use computational protein design to create IL-21 mimics with improved therapeutic properties.

## Computational protein design of an IL-21 mimic

IL-21 signals by inducing heterodimerization of the IL-21 receptor (IL-21R) with the common cytokine receptor 𝛾 chain (𝛾_c_), leading to the activation of JAK1 and JAK3, which are associated with the IL-21R and 𝛾_c_ intracellular domains, respectively, resulting in the phosphorylation and activation of STAT transcription factors^34^. Native IL-21 makes extensive interfaces with both receptor subunits (Fig. 1A) involving largely helical secondary structures, but the “backside” of IL-21 contains two long loops that we reasoned might reduce the stability of the molecule. Traditional protein engineering approaches that use directed evolution to identify small numbers of sequence changes are not able to replace extended structural segments such as these poorly ordered regions. Instead, we sought to construct mimics that retained the receptor-interacting interfaces, but with a more well-ordered and less protease-susceptible overall structure. As the structure of the full complex was not initially available, we began from the human IL-21/IL-21R (hIL-21/hIL-21R) complex structure^35^ and docked the human 𝛾_c_ (h𝛾_c_) from the human IL-2 complex structure^36^ on the hIL-21/hIL-21R complex structure. We then generated helical protein scaffolds with regions that superimpose perfectly on the receptor-interacting segments of native IL-21, but with improved inter-helix packing, short connecting loops, and minimal unstructured regions.

**Fig. 1.**
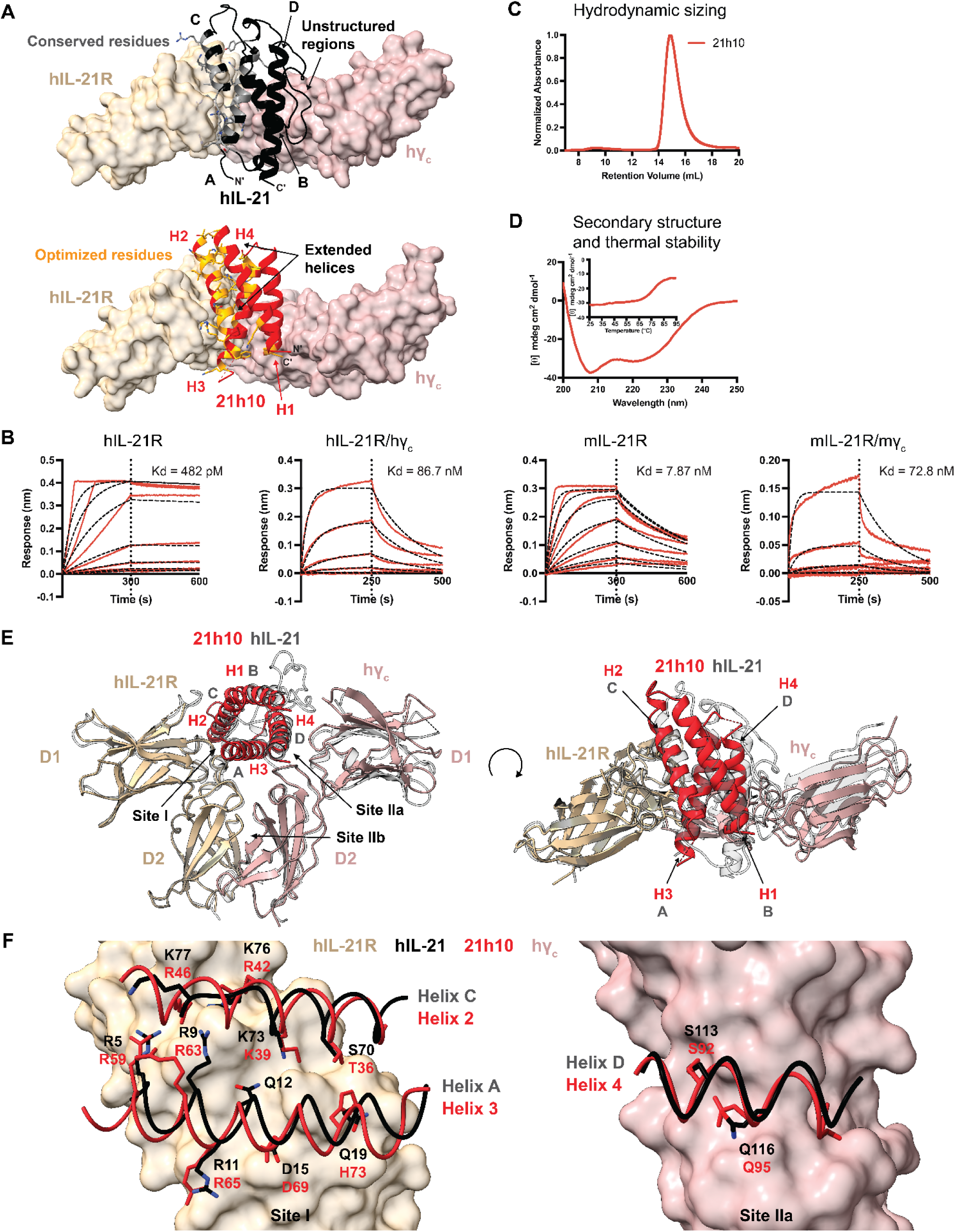
Computationally designed IL-21 mimic recapitulates receptor interactions with superior stability and human-mouse cross-reactivity. (A) The optimized IL-21 mimic, 21h10, is designed by recapitulating the helical bundle structure of the native hIL-21. Unstructured regions in the hIL-21 are removed, and short helices are extended to accommodate improved intramolecular packing. Some of the interface residues are conserved from the native hIL-21 for the design of initial hits. Other residues from the *de novo* scaffolds are further optimized, as shown. (B) Association and dissociation of 21h10 to human and murine IL-21R and 𝛾_c_ show concentration-dependent binding curves of 21h10. 21h10’s K_D_ against hIL-21R is 482 pM, hIL-21R/h𝛾_c_ is 86.7 nM, mIL-21R is 7.87 nM, and mIL-21R/m𝛾_c_ is 72.8 nM. For measuring K_D_ against hIL-21R and mIL-21R, IL-21R was immobilized on the biolayer interferometry (BLI) tip. For measuring K_D_ against hIL-21R/h𝛾_c_ and mIL-21R/m𝛾_c_, 𝛾_c_ was immobilized on the BLI tip and IL-21R was provided in solution with 1.5-fold molar excess to 21h10. Dotted lines show curve fits of raw data using a 1:1 Langmuir binding model. (C) The optimized IL-21 mimic, 21h10, exhibits monodispersed distribution when sized through size-exclusion chromatography using Superdex 75 10/300 GL column. (D) Wavelength scan from 200 nm to 250 nm confirmed 𝛼-helical secondary structure present in 21h10. Thermal melt from 25°C to 95°C with 222 nm scan revealed outstanding thermal stability of 21h10, with a T_m_ around 75°C. (E) Crystal structure of 21h10 in complex with hIL-21R and h𝛾_c_ (PDB: XXXX). (F) 21h10 conserves key molecular interactions in the site I interface (left) and site IIa interface (right) observed in the structure of the human IL-21 receptor complex. (Human IL-21 receptor complex, PDB ID: 8ENT).

Wild-type hIL-21 consists of four helices, with helix A (the first helix from the N-terminus) and helix C (the third helix) interacting with hIL-21R, and helix D (the fourth helix) interacting with h𝛾_c_ ^36,37^. Helix B (the second helix) and helix C are not long enough to form ideal intramolecular interactions and intermolecular contacts with the receptor chains, so we replaced the two long unstructured regions between helix A/helix B and helix C/helix D by extending helix B and helix C. We sampled different helical bundle “up-down-up-down” topologies by testing different connectivities between the helices; these differ from the “up-up-down-down” topology of native IL-21, and allow more ideal helical packing and loop geometries. Sequences were designed for these backbones in the context of hIL-21R using *Rosetta*, retaining the residues interacting with hIL-21R and anticipated to interact with h𝛾_c_. We stochastically generated designs with these properties using *Rosetta* helical sampling and loop-building methodology (see Methods), and selected for experimental characterization to validate their folding to the designed structure (fig. S1A).

## IL-21 mimic, 21h10, binds to human and murine IL-21 receptors

We obtained synthetic genes for the selected designs and assessed binding to hIL-21R using yeast surface display. Four designs bound hIL-21R, but none bound to h𝛾_c_ in the presence of hIL-21R (fig. S1B); this was not unexpected due to the unknown (at the time) h𝛾_c_ structure which was not incorporated in design stage. From a mutagenesis library around the tightest binder, 21d26, we identified a variant, 21JC15, that binds h𝛾_c_ weakly in an hIL-21R-dependent manner, as well as murine 𝛾_c_ (m𝛾_c_) in a murine IL-21R (mIL-21R)-dependent manner. Further interface optimization guided by site-saturation mutagenesis (SSM) and combinatorial library screening resulted in 21h10 with high affinity for both the human and murine IL-21 receptor complexes (Fig. 1B, and table S1). 21h10 is shorter than hIL-21 (102 vs. 131 amino acids) with sequence identity only in the conserved interface regions^38,39^. 21h10 was expressed at high levels in *Escherichia coli*, monodisperse in size-exclusion chromatography (Fig. 1C), and has a helical circular dichroism (CD) structure consistent with the design model. CD melting experiments revealed that 21h10 has high thermal stability, with a T_m_ around 75°C (Fig. 1D).

Copurification of 21h10 with the hIL-21R extracellular domain, followed by mixing this complex with the extracellular domain of h𝛾_c_ (see Methods, amino acids 33-232), resulted in the growth of crystals that diffracted anisotropically with resolution between 2.3 Å and 3.4 Å (fig. S2 and table S2). Solution of the structure by molecular replacement revealed the structure of 21h10 in complex with hIL-21R and h𝛾_c_ to be a Y-shaped receptor assembly similar to the wild-type hIL-21 complex^37^. 21h10 binds to hIL-21R at site I through helices 2 and 3, burying a surface area of 870 Å^2^, and it binds to h𝛾_c_ at site IIa through helix 4, burying a surface area of 423 Å^2^ (Fig. 1E and table S3). The key molecular contacts mediating receptor interaction are strongly conserved. At the site I interface, 21h10 interacts with hIL-21R through hydrogen bonds at residues K39, R46, R59, R63, and R65, and salt bridges at residues R42, R46, R53, and D69 (table S3). These side chains of 21h10 are nearly identically positioned to the corresponding side chains in hIL-21 despite different molecular topology (Fig. 1F). Similarly, at the site IIa interface, 21h10 binds to h𝛾_c_ through sparse hydrogen bonding at S92 and Q95 on helix 4 (table S3). These two residues are precisely conserved in helix D of hIL-21 (Fig. 1F), reflecting the ability of *de novo* protein design to reproduce important molecular interactions in a protein-protein interface accurately.

## 21h10 potently induces STAT phosphorylation and effector CD8+ T cell differentiation

Upon receptor binding, native IL-21 mediates downstream signaling by phosphorylating STAT1 and STAT3 in T cells^40^. While hIL-21 shows low cross-reactivity in murine cells (fig. S3A and table S4), 21h10 treatment *in vitro* led to phosphorylation of STAT1 and STAT3 with potency equivalent to native hIL-21 and mIL-21 in both human and murine CD8^+^ T cells, CD4^+^ T cells, and B cells (Fig. 2A, fig. S3B, and table S5).

**Fig. 2.**
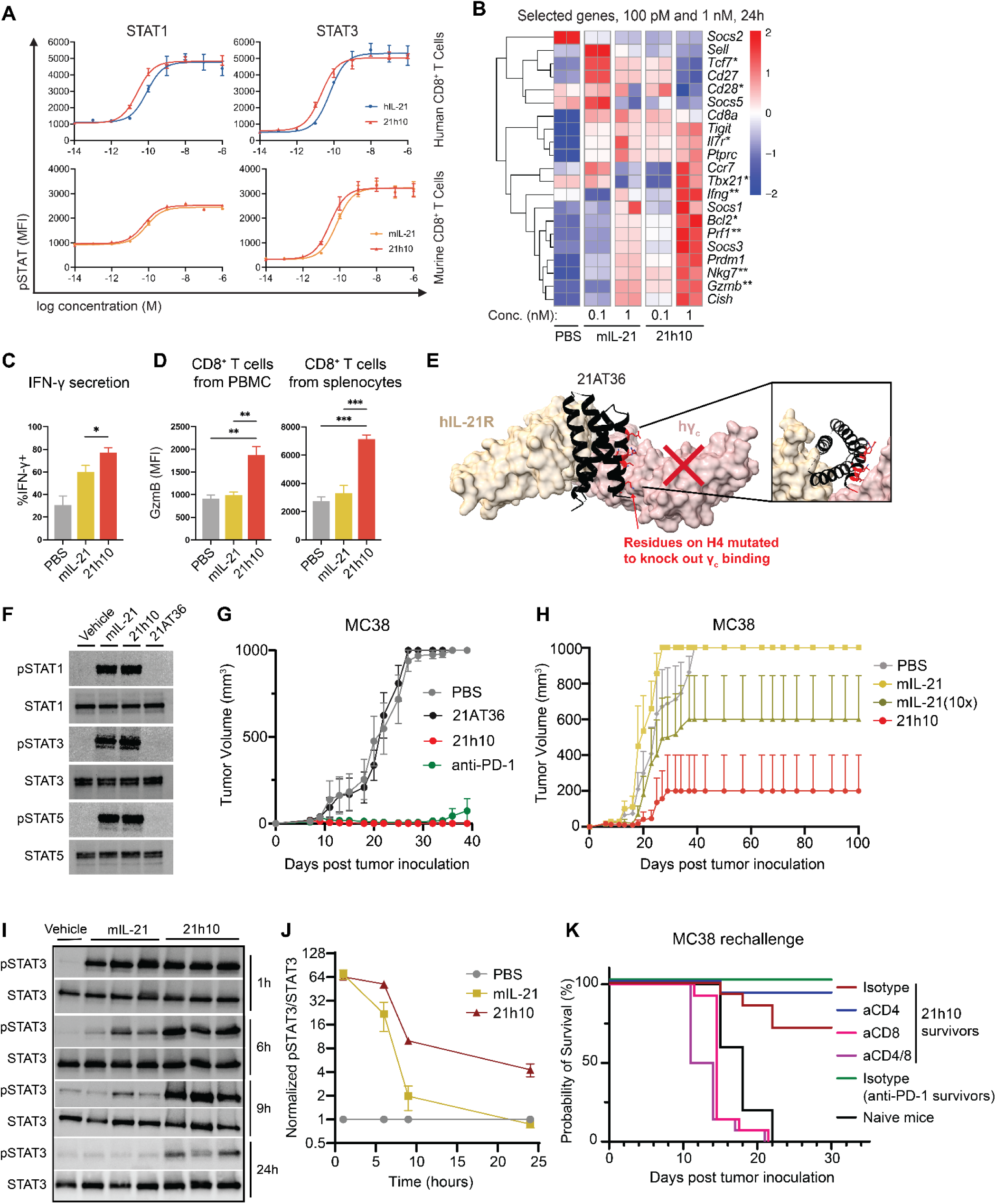
21h10 shows enhanced STAT signaling, gene expression, cellular phenotypes, and *in vivo* antitumor activity. (**A**) Pre-activated human and murine CD8^+^ T cells were treated with increasing doses of native IL-21 and 21h10 for 20 minutes and stained with AF488-phospho-STAT1 and AF647-phospho-STAT3 for flow cytometry analysis. (**B**) Genes related to cellular signaling and phenotype are compared for expression levels between murine CD8^+^ T cells treated with PBS, mIL-21, or 21h10 at 100 pM or 1 nM. * indicates memory-related genes, and ** indicates effector-related genes. Murine CD8^+^ T cells were pre-activated with TCR signals and treated with the cytokines at different concentrations for 24 hours, and cells were harvested for RNA-sequencing. (**C**) The percentage of IFN-𝛾-secreting cells within isolated CD8^+^ T cells upon treating PBS, mIL-21, or 21h10 at 1 nM, was quantified using flow cytometry after 5 hours of stimulation with PMA/Ionomycin in the presence of a protein transport inhibitor. (**D**) LCMV was inoculated into healthy mice along with daily injections of PBS, mIL-21 (2.5 µg per mouse), or 21h10 (2.5 µg per mouse). LCMV-specific CD8^+^ T cells from PBMC and spleen were analyzed for granzyme B expression. (**E**) Computational model of 21AT36, which is redesigned from 21h10 using ProteinMPNN, to bind IL-21R but not bind 𝛾_c_. The mutated residues are highlighted in red, which abolish interactions with 𝛾_c_. The 𝛾_c_ is included in the figure to show where the interface is positioned, not to imply that 21AT36 binds to 𝛾_c_. (**F**) Western blot for STAT and pSTAT with vehicle (PBS), mIL-21, 21h10, and 21AT36 in TRP1^high^ CD8^+^ T cells. (**G**) MC38-bearing mice were treated with PBS, 21AT36, 21h10, or anti-PD-1 for 14 days. (**H**) Mice were inoculated with MC38 and treated with PBS, mIL-21, 10-fold dose mIL-21, and 21h10. (**I**) Western blot for STAT and pSTAT in the spleens of mice treated with vehicle (PBS), mIL-21, and 21h10 (**J**) Normalized pSTAT3/STAT3 in spleens of treated mice over 24 hours from (**I**). (**K**) MC38 rechallenges with MC38-survivor mice from previous 21h10 or anti-PD-1 treatment.

Native IL-21 promotes effector CD8^+^ T cell differentiation in human CD8^+^ T cells *ex vivo*^3,41^ and plays a role in effector T cell (CD44^+^CD62L^-^) generation *in vivo*^42^. To investigate the effects of 21h10 on gene expression and effector CD8^+^ T cell differentiation, we cultured murine CD8^+^ T cells in media containing serum with 100 pM, 1 nM, 10 nM, and 100 nM of mIL-21 or 21h10 and analyzed gene expression by RNA-sequencing. 21h10 increased gene expression of effector molecules (*Tbx21*, *Prdm1*, *Prf1*, *Gzmb*) while reducing the expression of the genes associated with a resting state (*Tcf7*, *Sell*) as compared to mIL-21 at 100 pM and 1 nM. 100 pM of 21h10 elicited a similar gene expression profile as 1 nM of mIL-21 (Fig. 2B and fig. S3C); gene expression patterns were similar but not identical at higher doses (fig. S3D). 21h10 elicited similar or higher IL-21R and granzyme B expression levels in human and murine CD8^+^ T cells compared to hIL-21 and mIL-21, respectively (fig. S4A), and proliferation of murine CD8^+^ T cells at 1 nM and 100 pM of 21h10 was enhanced compared to mIL-21 (fig. S4, B to C). 21h10 promoted maximal differentiation of murine CD8^+^ T cells into CD44^+^CD62L^-^ effector T cells at 1 nM, while mIL-21 showed similar effect at 100 nM (fig. S4D) accompanied by a robust increase in the secretion of IFN-𝛾 at 1 nM (Fig. 2C). In the *in vivo* setting of mice infected with lymphocytic choriomeningitis virus (LCMV), 21h10 induced granzyme B expression in virus-specific CD8^+^ T cells, while mIL-21 had minimal effects (Fig. 2D); thus, 21h10 enhances effector CD8^+^ T cell differentiation both *in vitro* and *in vivo*.

## 21h10 has potent antitumor activity

Using ProteinMPNN^43^, we redesigned the 𝛾_c_-interface on helix H4 of 21h10 to abolish its receptor affinity, resulting in an antagonist, 21AT36, which binds to IL-21R but not 𝛾_c_ (Fig. 2E and fig. S5A). While mIL-21 and 21h10 induced phosphorylation of STAT1, STAT3, and STAT5, 21AT36 did not induce STAT phosphorylation (Fig. 2F and fig. S5B). *In vivo*, 21h10 led to rapid regression of MC38 adenocarcinoma, a highly immunogenic tumor model^44^, whereas 21AT36, as expected, did not show antitumor activity (Fig. 2G and fig. S6A).

21h10 exhibited enhanced antitumor activity in the MC38 model compared to native IL-21 despite similar activity in cell culture, outperforming both a molar equivalent and a 10-fold molar excess of native IL-21 (Fig. 2H). We hypothesized that this increased activity was due to the enhanced stability of 21h10, resulting in resistance to serum proteases, longer receptor occupancy time, and prolonged *in vivo* signaling. To test this, we treated mice with a single intravenous dose of native IL-21 or 21h10 and measured pSTAT3 in splenocytes over time. Despite similar initial levels of cytokine-induced pSTAT3, native IL-21-induced pSTAT3 decreased rapidly with time, whereas 21h10-induced pSTAT3 declined more slowly and remained detectable at 24 hours after injection (Fig. 2, I to J). 21h10 also retained greater *in vitro* activity after incubation with murine serum compared to native IL-21 (fig. S7A), consistent with the larger effects of 21h10 compared to native IL-21 during prolonged *in vitro* culture in serum-containing media (figs. S4 and S7B).

The improved *in vivo* activity of 21h10 was not due to off-target binding, as MC38 challenge in *Il21r*-/- mice showed no efficacy (fig. S6B). 21h10 treatment induced immune memory in MC38 treated mice, as rechallenged animals were protected. Memory responses were dependent on CD8^+^ T cells as protection was lost upon CD8^+^ and CD8^+^/CD4^+^ T cell depletion, but not in animals depleted of CD4^+^ T cells alone (Fig. 2K). Similarly, the antitumor response against MC38 is not dependent on B cells, as μMT mice which lack B cells showed responses to 21h10 that were equivalent to wild-type animals (fig. S6C). *In vitro*, tumor-targeting CAR-T cells generated in the presence of 21h10 exhibited enhanced cell expansion and cytotoxicity, with increased expression of CD25, granzyme B, and Bcl2 (fig. S8, A to C), a shift in cellular metabolism to a high metabolic state, and rapid elimination of MC38 adenocarcinoma when subjected to multiple rounds of tumor injection (fig. S8, D to F).

To determine the generalizability of our findings in MC38 adenocarcinoma, we assessed 21h10 in a less immunogenic tumor model. B16F10 is a model of poorly immunogenic melanoma known to respond to various immunotherapies, including cytokine therapies^32,33^. Responses to B16F10 can be augmented through the transfer of tumor-specific CD8^+^ T cells that recognize the TRP1 self-antigen with high or low affinity (TRP1^high^ and TRP1^low^ T cells, respectively), although these cells are insufficient to cause tumor regression on their own^45^. Transferring TRP1^high^ and TRP1^low^ T cells models an oligoclonal T cell response to a tumor-associated self-antigen, analogous to what is observed in immunogenic human melanoma. We adoptively transferred TRP1^high/low^ T cells and treated B16F10 melanoma-bearing mice with PBS, mIL-21, or 21h10. mIL-21 treatment delayed tumor growth compared to PBS; however, as we observed for MC38, 21h10 treatment exhibited more robust tumor regression, with several long-term surviving animals (Fig. 3A and fig. S9A). Moreover, 21h10 synergized with adoptive cell transfer of TRP1^high/low^ cells, as compared to no adoptive transfer, and 21h10 had stronger antitumor activity than was seen for the IL-2/IL-15 mimic Neo-2/15^32,33^; strikingly, 21h10 treatment combined with adoptive T cell transfer resulted in complete tumor regression in 2 out of 13 animals (Fig. 3, B to C, and fig. S9, B to C). We found similarly enhanced antitumor responses when 21h10 was combined with the anti-melanoma antibody TA99^32,33^ (fig. S10A). *Ex vivo* treatment of TRP1^high/low^ T cells with 21h10 prior to transfer into B16F10-bearing mice modestly decreased tumor growth (fig. S10B), suggesting that sustained signaling in the tumor microenvironment may be important for optimal antitumor ^46,47^ actions of 21h10.

**Fig. 3.**
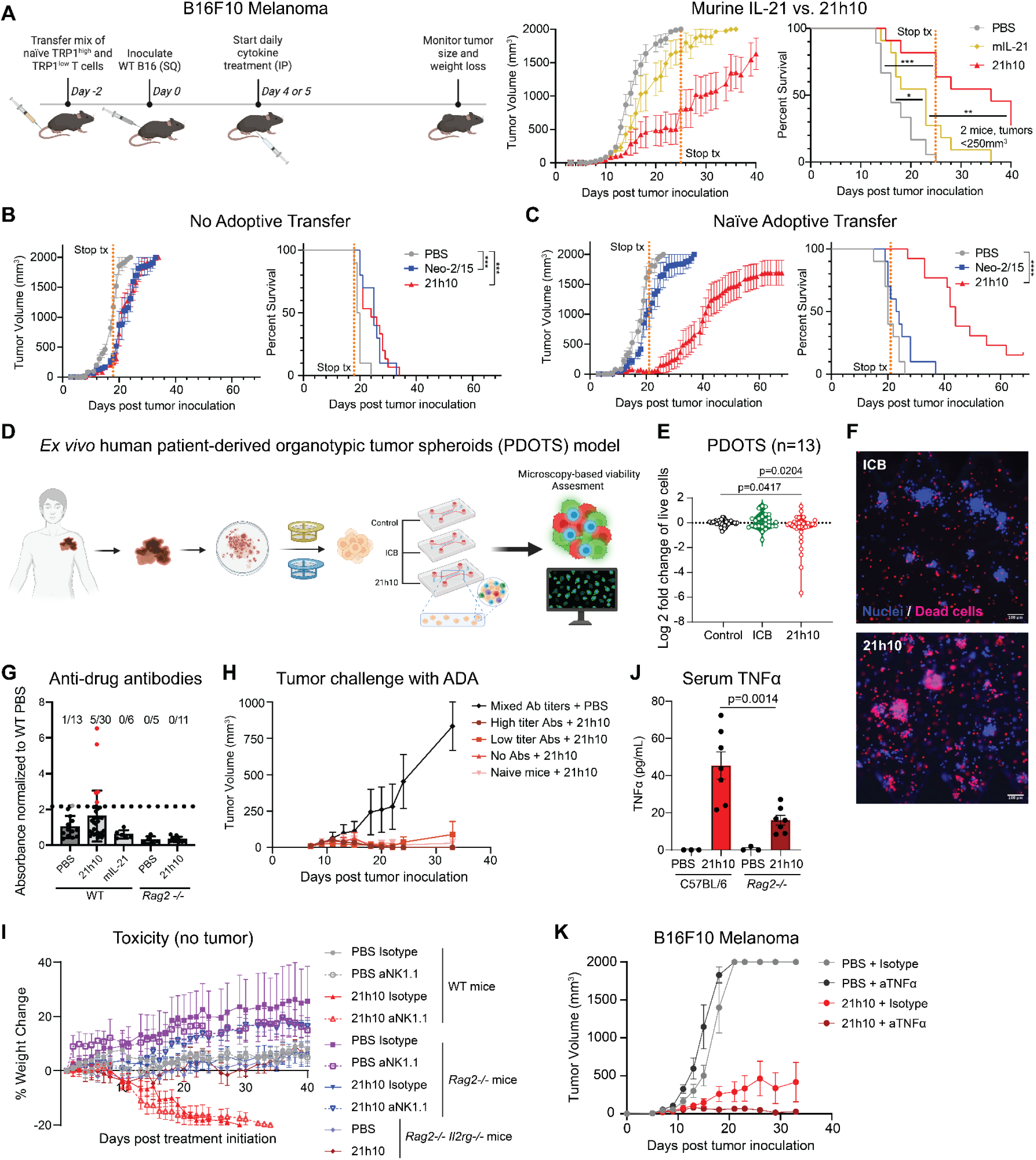
21h10 improves antitumor efficacy in *in vivo* B16F10 murine melanoma and *ex vivo* human PDOTs models, with non-neutralizing anti-drug antibodies, and toxicity alleviated with TNFα blockade. (**A**) Naïve CD8^+^ TRP1^high/low^ melanoma-specific T cells were adoptively transferred to mice prior to B16F10 inoculation. Cytokine therapy began on day 5 and continued every day until the stop of treatment (dashed line). (**B**) Mice received no prior T cell adoptive transfer before cytokine therapy. (**C**) Same as (**B**), but with adoptive transfer of naïve CD8^+^ TRP1^high/low^ T cells. (**D**) Scheme of PDOTS preparation. (**E**) Viability assessment of *ex vivo* human melanoma PDOTS following treatment of ICB (anti-PD-1, pembrolizumab, or anti-PD-1/anti-LAG-3), 21h10, or untreated control. (**F**) Representative images of PDOTS viability assessment shown in (**E**). PI-dead cells in red and Hoechst-nuclei in blue. (**G**) Mice died due to toxicity or after 40 days of treatment were bled and their serum was analyzed by ELISA for anti-drug antibodies against 21h10. Positivity (dotted line) was determined as two standard deviations above the mean of samples from mice treated only with PBS. Fractions indicate the number of mice classified as “positive” for anti-drug antibodies out of the total mice in each treatment group. (**H**) Mice with various titer levels of anti-21h10 antibodies were inoculated with MC38 tumors. Starting on day 6, mice were treated with 21h10 or PBS daily for 17 days. (**I**) Wild-type (WT), *Rag2-/-*, or *Rag2-/- Il2rg-/-* mice were treated with PBS or 21h10 daily and isotype or anti-NK1.1 depleting antibodies every three days. Mice were sacrificed when weight loss exceeded 20% of the initial starting weight. (**J**) Serum TNFα level was measured in C57BL/6 and *Rag2-/-* mice with PBS or 21h10 treatments. (**K**) Antitumor activity comparison on B16F10 melanoma between 21h10 and 21h10+aTNFα treatment groups.

To determine whether 21h10 had activity against human tumors, we used patient-derived organotypic tumor spheroids (PDOTS) model^48^. PDOTS, unlike organoids, are fresh tumor explants with intact immune compartment for *ex vivo* profiling of immunotherapy response of human tumors^48^. PDOTS were derived from patients with advanced melanoma, with the majority of samples from patients who progressed on standard of care checkpoint blockade (n=13, n=11 clinically resistant, table S6). 21h10 demonstrated activity against these resistant tumors, outperforming PD-1 blocking antibody (pembrolizumab) and a combinatorial immunotherapy of anti-PD-1 (nivolumab) and anti-LAG3 (relatlimab) antibodies (Fig. 3, D to F).

We next characterized the immunogenicity and tolerability of 21h10. Anti-drug antibodies were detected in 5 of 30 mice treated with 21h10 in antitumor experiments, with low-titer antibodies to 21h10 detectable at a 1:100 dilution at a concentration two standard deviations above the mean of the PBS controls. No anti-21h10 antibodies were detected in mIL-21-treated controls (Fig. 3G), consistent with the low sequence homology between native mIL-21 and 21h10. Anti-21h10 antibodies did not appear to neutralize 21h10. Mice vaccinated with multiple cycles of 21h10 to generate anti-drug antibodies were able to subsequently respond to 21h10 treatment in the MC38 model, showing no correlation with the level of anti-21h10 antibodies generated (Fig. 3H).

Although most animals could tolerate 21h10 at therapeutic doses, some animals exhibited weight loss, particularly at higher doses. This toxicity did not require NK cells, as it was not attenuated by treatment with NK cell-depleting antibodies (Fig. 3I and fig. S11A). In contrast, *Rag2-/-* mice, which lack T and B cells, were protected from 21h10 toxicity (Fig. 3I and fig. S11A), implicating adaptive immunity in 21h10 toxicity. We reasoned that cytokines produced by adaptive immune cells in response to 21h10 might play a role in toxicity. IFN-𝛾 was not required for toxicity, as treatment with 21h10 led to toxicity in *Ifng-/-* mice (fig. S11B). To evaluate whether a different 21h10-induced cytokine may be causing toxicity, we measured serum cytokines in 21h10-treated WT mice, *Rag2-/-*, and *Ifng-/-* mice. G-CSF, IL-17, and TNFα were all elevated in WT and *Ifng-/-* mice, but not in *Rag2-/-* controls (Fig. 3J and fig. S11C). When we co-administered TNFα blockade with 21h10, weight loss was mitigated, indicating that TNFα plays an important role in 21h10 toxicity (fig. S11D). In the B16F10 melanoma model, blockade of TNFα did not impair the antitumor activity of 21h10 (Fig. 3K).

## Mechanism of 21h10 potent antitumor activity

IL-21 has diverse roles within the immune system depending on the responding cells and the context in which it is produced^10^. To understand the mechanism of action of 21h10 *in vivo*, we performed single-cell RNA-sequencing (scRNAseq) on CD45^+^ enriched cells from the B16F10 melanoma, comparing mice treated with PBS, Neo-2/15, mIL-21, and 21h10 in addition to adoptively transferred TRP1^high/low^ CD8^+^ T cells. Using unbiased clustering, we identified populations of T cells, multiple monocyte/macrophage populations, dendritic cells, and smaller clusters of B cells, NK cells, granulocytes, melanoma cells, and fibroblasts (Fig. 4A and fig. S12A) in each treatment group (Fig. 4B). All T cell clusters were significantly expanded in mice treated with Neo-2/15 (Fig. 4, B to C), consistent with the known effects of IL-2 and IL-2 mimics on T cell proliferation and survival^49^, whereas 21h10, mIL-21, and PBS had modest effects on the relative proportion of T cells within the tumor microenvironment (Fig. 4, B to C). The T cell expansion observed with Neo-2/15 primarily came from CD8^+^ T cells, consistent preferential expansion of CD8^+^ T cells compared to CD4^+^ T cells by IL-2 ^50^ (Fig. 4D and fig. S13A). Like total T cells, TRP1-specific T cells, particularly low-affinity TRP1^low^ cells, expanded with Neo-2/15 (Fig. 4E). Although 21h10 treatment did not expand total CD8^+^ T cells, tumor-specific low-affinity TRP1^low^ cells accumulated in 21h10-treated tumors (Fig. 4E), consistent with studies showing that low-affinity T cell expansion can be greater than that of their high-affinity counterparts^51^. Native mIL-21 did not expand TRP1^high^ or TRP1^low^ T cells (Fig. 4E).

**Fig. 4.**
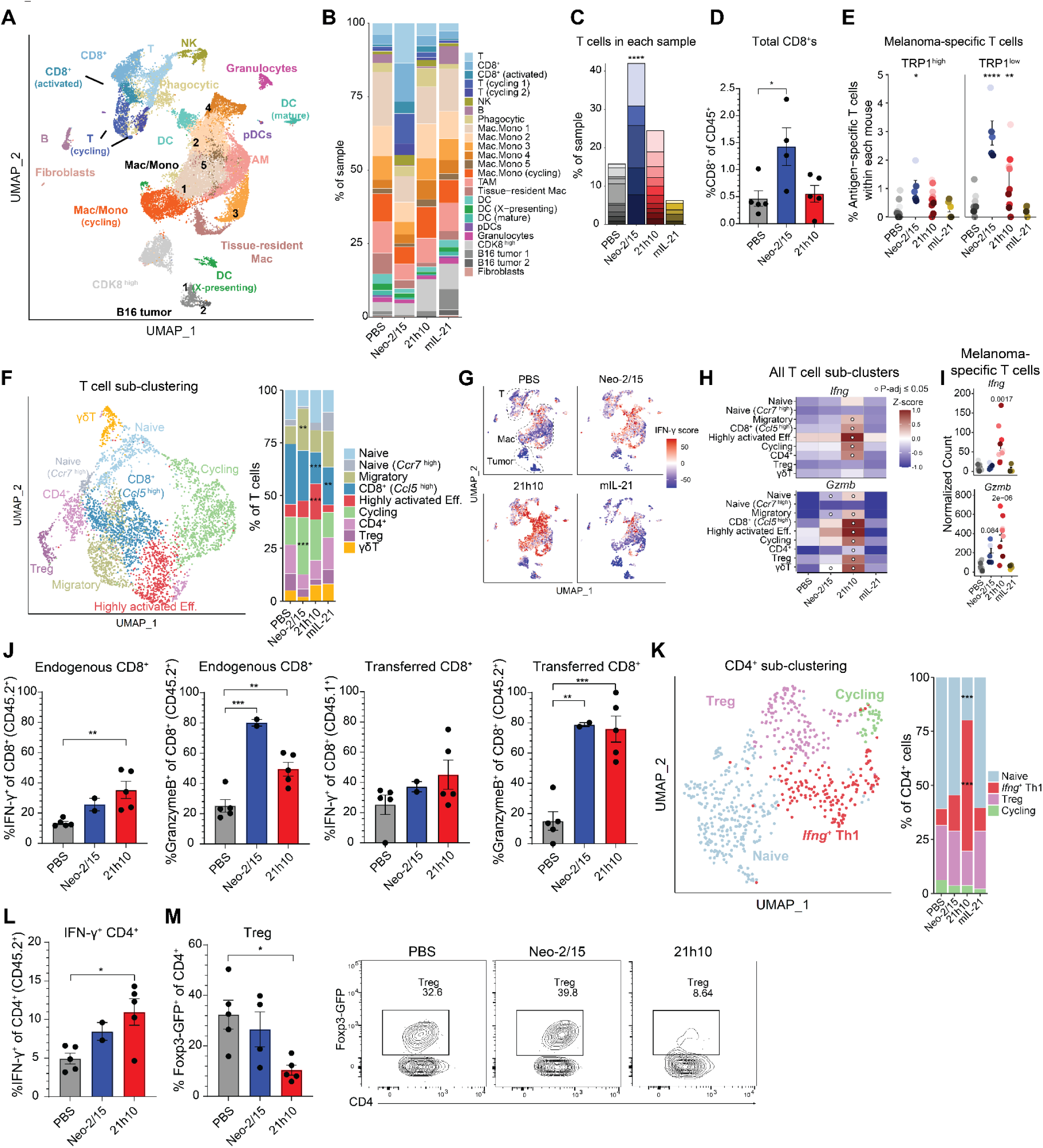
21h10 treatment results in expanded antitumor T cells with enhanced effector phenotype. (**A**) Mice were treated as in Fig. 3, A and C, but on day 15 of tumor growth, tumors were processed for single-cell RNA-sequencing. UMAP of all samples are combined. (**B**) scRNAseq cluster composition average across treatment groups. (**C**) T cells in each scRNAseq sample from (**B**). Different colors for each bar indicate individual mice from each group. (**D**) Flow cytometry quantification of CD8^+^ T cell infiltration. (**E**) TRP1^high^ and TRP1^low^ melanoma antigen-specific cells captured by sequencing. Two-way ANOVA with Dunnett’s multiple comparisons test vs. PBS groups. (**F**) T cells were sub-clustered from all samples from (**A**). The average T cell sub-cluster composition across treatment groups is shown on the right. Two-way ANOVA with Dunnett’s multiple comparisons test vs. PBS samples. (**G**) IFN-𝛾-response score total UMAP from (**A**) based on scaled and summed Hallmark IFN-𝛾-response genes for each cell. (**H**) Heatmap from pseudo-bulk differential gene expression analysis across T cell sub-clusters. Significant differences compared to PBS samples. (**I**) Similar to (**H**), but for TRP1 cells only, values from individual mice are shown. P-values displayed vs. PBS samples. (**J**) Flow cytometry quantification of CD8^+^ IFN-𝛾/granzyme B levels in endogenous or transferred CD8^+^ TRP1^high/low^ T cells. (**K**) CD4^+^ T cell sub-clustering UMAP and cluster composition. Two-way ANOVA with Dunnett’s multiple comparisons test vs. PBS samples. (**L**) Flow cytometry of IFN-𝛾^+^ CD4^+^ T cells. (**M**) Quantification and representative flow cytometry plots of Foxp3-GFP Treg cells. One-way ANOVA with Dunnett’s multiple comparisons for flow quantification and scRNA-seq total T cell comparison. For all dot plots with error bars: dots indicate individual mice, bars are S.E.M. *p ≤ 0.05, **p < 0.01; ***p < 0.001; ****p < 0.0001, or p-values displayed.

The high antitumor activity of 21h10 is striking, given the modest effect on T cell expansion. Several lines of evidence suggest that 21h10 enhances T cell effector functions. T cell sub-clustering identified two naïve T cell populations expressing *Tcf7*, a migratory cluster (*Itga4, Themis*), activated CD8^+^ T cells (*Ccl5, Gzmk*), highly activated CD8^+^ T cells expressing multiple immune checkpoint receptors (*Havcr2, Lag3*) and effector molecules (*Gzmb, Ifng*), a proliferative cluster (*Mki67, Birc5*) and CD4^+^ T cells (*Cd4, Cd40lg*), Tregs (*Foxp3, Ctla4*), and 𝛾𝛿 T cell clusters (Fig. 4F and fig. S12B). T cell expansion by Neo-2/15 mainly occurred in the migratory and cycling clusters, but 21h10 had greater effects on highly activated CD8^+^ T cells. No significant increase in T cell populations was observed with mIL-21. Neo-2/15 primarily affected pathways involved in cell growth and proliferation, while mIL-21 showed a similar, albeit weaker effect compared to PBS. 21h10 treatment was associated with IFN-𝛾/IFN-𝛼 response genes and growth pathways (fig. S12C) in all immune populations identified, including the myeloid and T cell clusters (Fig. 4G). 21h10 expanded the most activated CD8^+^ T cells and augmented *Ifng* and *Gzmb* gene expression in most T cell subclusters (Fig. 4H), including the transferred melanoma-specific CD8^+^ T cells (Fig. 4I), and correspondingly, IFN-𝛾 and granzyme B levels were increased in both endogenous and transferred T cells from 21h10-treated animals, as compared to PBS controls (Fig. 4J and fig. S13B).

There were fewer tumor-infiltrating CD4^+^ T cells than CD8^+^ T cells, as shown by both flow cytometry and scRNAseq. We subclustered the CD4^+^ T cells and identified four populations: naïve CD4^+^, cycling CD4^+^ T cells, Tregs, and Th1 cells expressing *Ifng* mRNA (Fig. 4K and fig. S12D). The *Ifng*^+^ Th1 cells preferentially expanded in response to 21h10 as compared to the other treatments, and had higher levels of IL-21R, analogous to what we observed with CD8^+^ T cells (Fig. 4, K to L, and figs. S13C and S14). Our scRNAseq analysis also suggested that 21h10 decreased Tregs, as we confirmed by flow cytometry using Foxp3-GFP reporter mice (Fig. 4M). Thus, 21h10 treatment robustly activates STAT1 and STAT3 in T cells in the tumor microenvironment, and this is accompanied by a shift toward highly activated, IFN-𝛾- and granzyme-B-producing, anti-melanoma CD8^+^ T cells as well as a relative increase in Th1 cells and decrease in Tregs. These findings help to explain the enhanced antitumor responses by 21h10 compared to mIL-21 or Neo-2/15.

## Discussion

Cytokine-based therapies for cancer have historically had limited efficacy and high toxicity^52–54^. Here, we show that a computationally-designed mimic of IL-21 with improved stability can have potent antitumor activity, both as a monotherapy and in combination with adoptive T cell therapies. 21h10 expanded highly activated effector CD8^+^ T cells in the tumor microenvironment, with the effect most pronounced for CD8^+^ T cells with known tumor reactivity. In contrast, the IL-2/IL-15 mimic Neo-2/15, expanded multiple T cell populations without a preferential effect on the differentiation of effector cells, providing a mechanistic explanation for the difference in antitumor activity between these two cytokine mimics. Antitumor T cell responses can display a range of receptor affinities, with low-affinity responses typically seen when a self-antigen is targeted^55,56^. Using single-cell RNA sequencing, we found that CD8^+^ T cells recognizing the melanoma self-antigen TRP1 with either high or low affinity were both augmented by 21h10, but strikingly, low-affinity TRP1-specific T cells were preferentially increased, indicating that 21h10 selectively amplified lower affinity antitumor T cells. By contrast, PD-1 blockade immunotherapies appear to act on high-affinity neoantigen-reactive clonotypes^57^, and immunotherapies that can augment a broader spectrum of TCR affinities are currently lacking. Low-affinity T cells often fail to receive a signal strong enough to overcome the TCR signaling threshold^58^, and thus are frequently unable to contribute to antitumor immunity^45^ – by expanding the TRP1^low^ population, 21h10 broadens the repertoire of the antitumor immune response, potentially explaining the efficacy observed in *ex vivo* cultures of immunotherapy resistant human melanoma. 21h10 also decreased the Treg population while inducing an effector phenotype on CD8^+^ T cells. Although IL-21 has important roles in B cell differentiation, class switching, and NK cell activation, neither B nor NK cells were significantly altered in our *in vivo* tumor models.

Although we chose 21h10 for its ability to mimic IL-21 signaling *in vitro*, the resulting *de novo* protein showed dramatically superior performance *in vivo*. We showed that 21h10 has increased serum stability compared to native mIL-21, leading to sustained potency *in vivo*. These properties increase the duration of signaling *in vivo*, leading to enhanced phenotypic changes and, ultimately, improved antitumor responses. These improvements highlight the robustness of *de novo* protein design approaches for developing therapeutic candidates with improved properties. The full human/mouse cross-reactivity and engineerability of 21h10 should make it straightforward to generate targeted and conditionally active versions^33^ to mitigate toxicity and target activity to the tumor microenvironment.

In summary, we have developed a potent IL-21 mimic with activity on both human and mouse cells that exhibits robust antitumor activity in multiple tumor models. 21h10 has antitumor activity as a monotherapy, synergizes with adoptive cell therapy in B16F10 melanoma model, and is curative in a highly immunogenic MC38 adenocarcinoma model. Activity of 21h10 in human refractory melanoma PDOTS and its superiority on clinically approved immune checkpoint blockade (ICB) treatments suggest the potential for rapid clinical translation to treat tumors unresponsive to checkpoint blockade and other existing therapies.

## Methods

### Computational design of *de novo* IL-21 mimic

The crystal structure of hIL-21 with hIL-21R (PDB: 3TGX) was used to design the mimics of native IL-21. The *PyRosetta* script with PDBInfoLabel metadata implementation generated IL-21-like scaffolds^32^ (Appendix A). The following residues from the human IL-21 are designated to be fixed during scaffold generation and interface residue design: R2, I5, R6, R8, Q9, L10, I11, D12, I13, D15, Q16, K18, Y20, R62, I63, V66, S67, K69, K70, R73, K74, P75, P76, S77, K98, E99, E102, R103, K105, S106, Q109, K110, H113, and L116 (Appendix B). The non-fixed residues are designed using *Rosetta* FastDesign and relaxed using *Rosetta* FastRelax with the ‘beta_nov16’ score function. 185 designs were filtered with *Rosetta* score metrics: packstat>0.6, score_per_residue<-2.3, and sspred>0.8. Fast forward folding was used on the designs to filter them based on their probability of folding, giving 36 designs to be tested by experimental screening.

### Yeast display and fluorescent-activated cell sorting

The amino acid sequences of the designed proteins were reverse-translated to DNA sequences based on the codon frequency table of *Saccharomyces cerevisiae,* along with pETcon3 vector-homologous sequences appended to both 5’ and 3’ termini. Using competent yeast cells, *EBY100*, the DNA gene blocks of the designs were cloned into linearized pETcon3 vectors by yeast homologous recombination. The vector was linearized by 100-fold overdigestion by NdeI and XhoI (New England Biolabs) and then purified by gel extraction (Qiagen). Yeast transformations are validated by colony sequencing; validated colonies are grown in C-Trp-Ura medium and induced in SGCAA medium as previously described^59^. SGCAA induced the designed proteins to be expressed and presented on the yeast surface through Aga2p membrane proteins. The induced cells with the designed proteins on their surfaces were labeled with two fluorophores - FITC and PE. Anti-c-myc mAb conjugated to FITC was used to label the myc tag to represent the degree of protein expression. For IL-21R labeling, either human or murine IL-21R with human Fc-tag (R&D Systems Cat# 991-R2, R&D Systems Cat# 596-MR) were used, and for receptor complex labeling, either human IL-21R with His-tag (R&D Systems Cat# 9249-R2) and human 𝛾_c_ with Fc-tag (Acro Biosystems Cat# ILG-H5256), or mIL-21R with His-tag (Sino Biological Cat# 51184-M08H) and murine 𝛾_c_ with Fc-tag (R&D Systems Cat# 784-MR) were used. The target receptor fused with the Fc domain is labeled by the biotinylated ZZ domain of protein A and streptavidin-PE (SA-PE), which can bind to the biotin of protein A. PE represents the level of receptor binding. For binding hits, their signal is proportional to the protein expression level. Yeast was analyzed by flow cytometers (BD Accuri C6, Thermo Fisher Attune NxT) or sorted by a fluorescence-activated cell sorting (FACS) cell sorter (Sony SH800).

### Mutagenesis and affinity maturation

Using a random mutagenesis library kit (Agilent Cat# 200550), the 21d26 DNA was amplified, and the library was partially sequenced using BL21(DE3) (New England Biolabs). The sequencing showed that approximately 2-4 mutations were applied to the design while the DNA was amplified with the error-prone PCR. Next, the library was transformed to yeast using electroporation with an approximated diversity of 3.4E7. The yeast library was sorted for direct evolution using FACS with different combinations and concentrations of labeled receptor subunits with the abovementioned labeling. With three consecutive sorts with either human receptor complex labeling (1 µM of unlabeled hIL-21R and 1 µM of labeled h𝛾_c_) saturated sorted library with mutants (including 21JC15) with h𝛾_c_-binding capability.

To analyze and enhance the interface affinities of the interfaces of the 21JC15, we used SSM to analyze the two interfaces of the protein in detail. The protein sequence of interest was split into halves with corresponding single mutations and synthesized as two fragments for assembly (Integrated DNA Technologies). Assembly was done with PCR with pETcon3-specific 5’ and 3’ primers. Sortings with four different receptor conditions – (1) labeled hIL-21R, (2) unlabeled hIL-21R with labeled h𝛾_c_, (3) labeled mIL-21R, and (4) unlabeled mIL-21R with labeled m𝛾_c_ – saturated mutants with mutants with favorable mutations, and depleted mutants with unfavorable mutations. DNA of the sorted SSM library was prepared prior to MiSeq (Illumina Cat# MS-102-3001) using Zymoprep (Zymo Research Cat# D2001), qPCR, and gel extraction (Qiagen). The sorted SSM library revealed several mutations that can improve binding affinities to each receptor - hIL-21R, h𝛾_c_, mIL-21R, and m𝛾_c_.

By merging the positive mutations, we built a combinatorial library that incorporated the mutations by using a PCR-based assembly of pools of 8 sets of primers (Integrated DNA Technologies) incorporating mutations with degenerate codons, which was then sorted against human or murine receptors. Sorting it yielded multiple candidates with significantly improved affinities to the receptors. The K_D_ of the candidates toward each of the receptors was measured by biolayer interferometry (BLI) (ForteBio) (See Methods).

### Protein expression and endotoxin removal

The DNA fragments (Integrated DNA Technologies) that encode the designed proteins were cloned into pET-29b(+) plasmid with an N-terminal polyhistidine tag. The cloned plasmids were transformed into competent BL21 cells (New England Biolabs), whose cultures were grown in Terrific Broth II and induced using 1 mM isopropyl b-D-thiogalactopyranoside (IPTG). The cultures were grown in baffled flasks at 37°C within 225 rpm shakers until they reached approximately OD_A600_=0.8 for induction. The cultures were induced for 14 hours at 18C. The harvested cultures were lysed by sonication (Qsonica Q500). The lysed cultures were ultracentrifuged at 18,000g for 30 minutes and purified using immobilized metal affinity chromatography (IMAC). The elution proteins were separated by HPLC size-exclusion chromatography using Superdex 75 10/300 GL column (GE Healthcare) on Akta (GE Healthcare AKTA Pure). Endotoxins in protein samples from *Escherichia coli* cultures were removed by loading the SEC-purified protein onto the Ni-NTA column and washing with endo-removal IMAC wash buffer (TBS, 1% CHAPS), which were then eluted, and the imidazole was removed by dialysis. The endotoxin levels were measured using Endosafe LAL cartridge with 0.1 EU/mL sensitivity (Charles River PTS201F) on Endosafe machine (Charles River NexGen-MCS).

### Protein expression of 21h10 receptor complex for crystallography

21h10 was expressed in *Escherichia coli* with a polyhistidine tag on its N-terminus and TEV protease cleavage site in between. After expression, the polyhistidine tag was removed using polyhistidine-tagged TEV protease and re-purified by running the lysate through Ni-NTA resin to remove TEV protease and size exclusion chromatography to purify the 21h10. The protein was then concentrated using a 5 kDa MWCO Amicon concentrator. The hIL-21R extracellular domain with glycan-reduced mutations (N78Q/N85Q/N106D/N116Q) was cloned into the pAcGP67a baculovirus vector with a C-terminal 6xHis-tag, and baculovirus was prepared by co-transfection of BestBac^TM^ DNA (Expression Systems) and pAcGP67a DNA into *Spodoptera frugiperda* (Sf9) cells. The virus harvested from Sf9 cells was used to infect *Trichoplusia ni* (Hi5) cells. The supernatant was harvested from Hi5 cells 72 hours after infection and incubated together with 21h10 on Ni-NTA resin (Qiagen) for 3 hours prior to elution with 0.3 M imidazole followed by purification using size-exclusion chromatography (SEC) on a Superdex 200 column (GE Healthcare). Finally, the h𝛾_c_ extracellular domain with an N-terminal truncation of 32 residues was cloned into pAcGP67a for baculovirus generation, and Hi5 cells were infected in culture media containing 5 µM kifunensine. The supernatant was harvested from Hi5 cells 72 hours after infection, deglycosylated by overnight incubation with Endo Hf (New England Biolabs), and purified by SEC on a Superdex 200 column.

### Biolayer interferometry (BLI)

Binding data were collected using OctetRED96 and OctetR8 (Sartorius). Octet binding buffer (HEPES+1% w/v Bovine Serum Albumin) was used for all reactions. To measure the binding affinity to hIL-21R or mIL-21R, biotinylated hIL-21R (R&D Systems Cat# AVI9249) or biotinylated mIL-21R (Acro Biosystems Cat# ILR-M82E3), respectively, were immobilized on streptavidin-coated biosensors (SAForteBio) at 5 μg/mL in the octet binding buffer, with 2-fold serial dilutions of the ligand starting at 100 nM, with 300 seconds for association and another 300 seconds for dissociation. Similarly, to measure binding affinity of 21AT36 to hIL-21R or mIL-21R, biotinylated hIL-21R (R&D Systems Cat# AVI9249) or biotinylated mIL-21R (Acro Biosystems Cat# ILR-M82E3), respectively, were immobilized on streptavidin-coated biosensors (SAForteBio) at 5 μg/mL in the octet binding buffer, with 3-fold serial dilutions of the ligand starting at 100 nM, with 300 seconds for association and another 300 seconds for dissociation.

To measure the binding affinity to hIL-21R/h𝛾_c_ or mIL-21R/m𝛾_c_, biotinylated h𝛾_c_ (Acro Biosystems Cat# ILG-H85E8) or biotinylated m𝛾_c_ (Acro Biosystems Cat# ILA-M82E3), respectively, were immobilized on streptavidin-coated biosensors (SAForteBio) at 5 μg/mL in the octet binding buffer and tested with 3-fold serial dilutions of the ligand starting at 500 nM, for 300-second association and 300-second dissociation in the presence of 1.5-fold molar excess of corresponding hIL-21R (R&D Systems Cat# 9249-R2-100) or mIL-21R (Sino Biological Cat# 51184-M08H). For ligands, hIL-21 (R&D Systems Cat# 8879-IL), mIL-21 (R&D Systems Cat# 594-ML), and 21h10 (see Methods) were used. Data was processed using ForteBio Data Analysis Software version 9.0.0.10. and the parameters were reported with standard error. All data are summarized in table S1.

### Circular dichroism and thermal stability assay

Far-ultraviolet circular dichroism measurements were performed with a spectropolarimeter (JASCO J-1500). The proteins were dissolved in PBS (pH 7.4) within 1 mm path length cuvettes at 0.5 mg/mL. The wavelength scan was performed within the 195 nm to 260 nm wavelength range. The thermal melt experiment was performed by heating samples (step 2°C/minute) from 25°C to 95°C and cooled back down to 25°C while observing absorption at 222 nm.

### Crystallization, data collection, and refinement

Deglycosylated 𝛾_c_ was mixed at an equimolar ratio to the co-purified IL-21/IL-21R complex. The ternary complex was concentrated to 10 mg/mL and treated with carboxypeptidase A and B. Crystals of the IL-21 ternary complex were grown by sitting drop vapor diffusion with 100 nL of the complex, mixed with an equal volume of 0.2 M potassium thiocyanate, 0.1 M Bis-Tris propane pH 6.5, 20% w/v PEG-3350, and crystals were harvested and cryoprotected using 30% ethylene glycol.

Diffraction data were collected at Stanford Synchrotron Radiation Laboratory beamline 12-2. Data were indexed, integrated, and scaled to 2.3Å resolution using XDS^60^. Due to anisotropy in the crystal diffraction, a pseudoellipsoidal diffraction limit was applied using Staraniso^61^. The structure was solved by molecular replacement in Phaser^62^ using models derived from structures of the IL-21/IL-21R binary complex (PDB: 3TGX)^35^ and a 𝛾_c_/nanobody complex (PDB: 7S2R)^63^, identifying four copies of 21h10, four copies of IL-21R, and two copies of 𝛾_c_ in the asymmetric unit. The structure was completed by iterative cycles of rebuilding and refinement in Coot^64^ and Phenix^65^. Noncrystallographic symmetry restraints were used in refinement. Data collection and refinement statistics are presented in table S2. The crystal structure has been deposited in the RCSB protein data bank with accession code PDB: XXXX. All crystallographic software was installed and configured by SBGrid^66^. Analysis of interfaces was conducted using PISA^67^ (table S3). Structure-related figures were generated using ChimeraX^68^ (UCSF).

### STAT phosphorylation assay in T and B cells

All experiments involving mice at NHLBI were performed using protocols approved by the NHLBI Animal Care and Use Committee and followed National Institutes of Health (NIH) guidelines for the use of animals in intramural research. Murine CD4^+^ and CD8^+^ T cells were isolated from the spleens of 6- to 8-week-old C57BL/6 mice through negative selection and magnetic separation (Stem Cell Technologies Cat# 19852 and Cat# 19853). Purified murine CD4^+^ and CD8^+^ T cells were pre-activated with plate-bound anti-mouse CD3 (2 µg/mL; clone 145-2C11, BioXCell Cat# BE0001-1) and soluble anti-mouse CD28 (1 µg/mL; clone 37.51, BioXCell Cat# BE0015-1) and cultured in complete RPMI media for 48 hours at 37°C. Human CD4^+^ and CD8^+^ T cells were isolated from the buffy coats of healthy human volunteers obtained from NIH Blood Bank. Buffy coats are produced as a by-product of a volunteer whole blood donation for transfusion. The buffy coats are separated from the packed red cells and plasma component during the process of manufacturing the red cells and plasma for transfusion. The buffy coat units are thus already in existence (pre-existing material) and would otherwise be discarded if not distributed for research use. Lastly, they are irreversibly anonymized prior to distribution. These characteristics meet the FDA 45 CFR46 2018 Revisions of the Common Rule, which state that pre-existing, otherwise discarded, anonymized human cells/tissue are exempt from the requirement for IRB review and informed consent. Peripheral blood mononuclear cells (PBMCs) were isolated using a lymphocyte separation medium based on density gradient centrifugation. CD4^+^ and CD8^+^ T cells were then purified by negative selection and magnetic separation (Stem Cell Technologies Cat# 17952 and Cat# 17953). Purified human CD4^+^ and CD8^+^ T cells were activated by plate-bound anti-human CD3 (2 µg/mL; clone UCHT1 (Leu-4) (T3), BioXCell Cat# BE0231) and soluble anti-human CD28 (1 µg/mL; clone 9.3, BioXCell Cat# BE0248) and cultured in complete RPMI medium for 48 hours at 37°C.

Murine B cells were isolated from the spleens of 6- to 8-week-old C57BL/6 mice through negative selection and magnetic separation (Stem Cell Technologies Cat# 19854). Purified murine B cells were pre-activated with anti-mouse CD40 (clone FGK45, BioLegend Cat# 157503) for 48 hours at 37°C in complete RPMI media. Human B cells were isolated from the buffy coats from healthy human volunteers obtained from NIH Blood Bank after written informed consent in accordance with the approval of the institutional human ethics committee. Peripheral blood mononuclear cells (PBMCs) were isolated using a lymphocyte separation medium based on density gradient centrifugation. Human B cells were then purified by negative selection and magnetic separation (Stem Cell Technologies Cat# 17954). Purified human B cells were pre-activated with anti-human CD40 (clone HB14, BioLegend Cat# 313020) for 48 hours at 37°C in complete RPMI media.

After 48 hours of culture, the pre-activated cells were rested overnight in a complete medium and restimulated with native IL-21 and IL-21 mimics at different concentrations (from 10 fM to 1 µM) for 20 minutes, respectively. For testing the heat stability, mIL-21, hIL-21, and 21h10 were pre-incubated overnight at 37°C in serum before treating the cells. Cells were then fixed, permeabilized, and stained with AF488-phosphoSTAT1 (1:100, BD Biosciences Cat# 612596) and AF647-phosphoSTAT3 (1:100, BD Biosciences Cat# 651008) for flow cytometry. Cells were analyzed using a BD LSR Fortessa X-20 Cell Analyzer flow cytometer (BD Biosciences).

### *In vitro* CD8^+^ T cell culture for flow cytometry

Purified murine and human CD8^+^ T cells were cultured in the presence of anti-mouse CD3 and CD28 or anti-human CD3 and CD28, respectively (as described above), together with mIL-21 or hIL-21 and 21h10 at 1, 10, 100 and 1000 ng/mL concentrations in complete RPMI medium for 72 hours at 37°C. Cells were harvested and stained with either APC anti-mouse IL-21R (1:100, Biolegend, Cat#131910) or APC anti-human IL-21R (1:100, Biolegend, Cat#347808) and BV421 anti-mouse/human granzyme B (1:100, Biolegend, Cat#396414) for flow cytometry. Cells were analyzed using a BD LSR Fortessa X-20 Cell Analyzer flow cytometer (BD Biosciences).

### Murine CD8^+^ T cell proliferation assay

For *in vitro* proliferation experiment, CD8^+^ T cells isolated from TRP1^high^ mice using EasySep Mouse CD8^+^ T Cell Isolation Kit (StemCell Cat# 19853) were stained with Cell Trace Violet (Invitrogen Cat# C34557) at a 1:1000 dilution in PBS and plated at 50,000 cells/well in a 96-well plate. Cells were plated with 3 µL/million cells of Dynabeads Mouse T-Activator CD3/CD28 (Gibco Cat# 11453D) along with various concentrations (0.001-10 nM) of mIL-21 (R&D Systems Cat# 594-ML-100/CF) or 21h10 in RPMI complete. Cytokines were incubated prior to addition to the cells for 24 hours at 37°C in healthy human serum. After 72 hours of treatment, cells were collected and analyzed by flow cytometry for CTV-labeled populations along with the following: APC anti-CD8a (clone 53-6.7, BioLegend Cat# 100712), Brilliant Violet 711 anti-CD44 (clone IM7, BioLegend Cat# 103057), PE anti-CD69 (clone H1.2F3, BioLegend Cat# 104508), and Zombie NIR (BioLegend Cat# 423105). Zombie NIR was used at a 1:500 dilution and antibodies were used at a 1:100 dilution.

### RNA-seq sample preparation

Murine CD8^+^ T cells were isolated from the spleens of 6- to 8-week-old C57BL/6 mice through negative selection and magnetic separation (Stem Cell Technologies Cat# 19853). Purified murine CD8^+^ T cells were pre-activated with plate-bound anti-mouse CD3 (2 µg/mL; clone 145-2C11, BioXCell Cat# BE0001-1) and soluble anti-mouse CD28 (1 µg/mL; clone 37.51, BioXCell Cat# BE0015-1) and cultured in complete RPMI media for 48 hours at 37°C. Cells were rested overnight and restimulated with mIL-21 or 21h10 at 100 pM, 1 nM, 10 nM, and 100 nM concentrations for 24 hours in complete RPMI media at 37°C. Cells were then lysed in TRIzol reagent (Invitrogen Cat# 15596026), RNA was isolated using Direct-zol RNA MiniPrep Kit (Zymo Research Cat# R2052), and the libraries were prepared for RNA-Sequencing using KAPA mRNA HyperPrep Kit (Roche Cat# KK8440) that was performed in the NHLBI DNA sequencing core.

### RNA-seq sample analysis

The libraries were barcoded (indexed) and sequenced on an Illumina NovaSeq platform. Sequenced reads were obtained with the Illumina CASAVA pipeline and mapped to the mouse genome (mm10/GRCm38) using TopHat 2.0.11. Raw counts that fell on exons of each gene were calculated and normalized by using RPKM (Reads Per Kilobase per Million mapped reads). Differentially expressed genes were identified with the R Bioconductor package “edgeR”, and expression heat maps were generated with the R package “pheatmap”.

### Murine T cell differentiation assay

All mice underwent experiments following guidance from the University of Washington Institutional Animal Care and Use Committee, protocol number 4397-01. Mice used for experiments harbored a non-perturbing Yellow Fluorescent Protein (YFP) reporter at the endogenous *Tcf7* locus as previously described^69^ and were donated by Avinash Bhandoola. Spleens were harvested from mice, and a single-cell suspension was generated by massaging spleens between rough glass slides and filtering over 50 µm mesh. Red blood cells (RBC) were lysed in RBC lysis buffer (150 mM NH_4_CL, 10 mM NaHCO_3_, 1 mM EDTA) for 3-5 minutes at room temperature. Splenocytes were incubated in Fc blocking buffer (10% 2.4G2 cell supernatant in FACS buffer, 0.5% BSA in HBSS) prior to CD8^+^ T cell isolation. CD8^+^ T cells were isolated using the CD8a^+^ T Cell isolation kit (Miltenyi Cat# 130-095-236) following the manufacturer’s instructions. CD8^+^ T cells were counted, stained with 5 µM CellTrace Violet (ThermoFisher) for 20 minutes at room temperature, and quenched with RPMI medium with 10% Fetal Bovine Serum (VWR) prior to activation.

One day prior to activation, tissue culture plates were coated with 1 µg/mL anti-CD3ε (BioLegend, Ultra-LEAF, Clone 145-2C11) and 0.5 µg/mL anti-CD28 (CyTek *in vivo* ready, Clone 37.51) in PBS and incubated overnight at 4°C. On the day of activation, plates were allowed to reach room temperature, washed 2x with room temperature PBS, and CTV-labeled T cells were resuspended in T cell medium (TCM, 85% RPMI 1640 with L-glutamine, 10% heat-inactivated Fetal Bovine Serum, 1x Pen-Strep-Glutamine, 20 mM HEPES, 1 mM Sodium Pyruvate, 0.1 mM Non Essential Amino Acids, and 50 µM β-mercaptoethanol (BME)) supplemented with 100 U/mL recombinant human IL-2 (Peprotech Cat# 200-02). T cells were seeded at 3×10^5^ cells per well of a pre-coated 24-well culture plate in 1.2 mL of TCM. Cells were incubated at 37°C 5% CO_2_.

After 48 hours, T cells were pooled, counted, centrifuged, and resuspended at 2.5×10^5^ cells/mL in a 24- or 96-well cell culture plate in TCM at 200 µL/well in uncoated 96-well plates and 1.2 mL/well in 24 well plates. TCM was supplemented as indicated in each figure either with PBS, mIL-21 (R&D Systems Cat# 594-ML), or 21h10 diluted in PBS carrier to a final concentration of 0.01 nM, 1 nM, and 100 nM. Cells were incubated at 37°C 5% CO_2_ for an additional 48 hours and collected directly for extracellular flow cytometry analysis or counted and re-stimulated for intracellular cytokine staining (ICS).

For ICS, cells were counted and resuspended in round bottom 96 well plates at 2×10^5^ cells per 100 µL of 1× Cell Stimulation Cocktail (eBioscience 500× stock) and incubated at 37°C 5% CO_2_ for 1 hour. Cells were then supplemented with Protein Transport Inhibitor (eBioscience 500x stock) to a final concentration of 1x and incubated at 37°C 5% CO_2_ for an additional 4 hours. Cells were then stained with 1:100 dilution Zombie Near IR (BioLegend Cat# 423105) for 15 minutes at room temperature, centrifuged and resuspended in the Fc blocking buffer, and incubated for 15 minutes on ice. Cells were then fixed and permeabilized for 15 minutes on ice using 100 µL/well of Cytofix/Cytoperm (BD), washed 2x with 1x Perm/Wash Buffer (BD), and incubated with anti-IFN-𝛾 (BioLegend, clone XMG1.2, PE) at 1:100 in 1x Perm/Wash Buffer for 30 minutes on ice. Cells were washed by an additional 2x with 1x Perm/Wash buffer and resuspended in FACS buffer for analysis on the Attune NxT (ThermoFisher).

For extracellular flow staining, cells were transferred to 96-well round bottom plates, centrifuged, resuspended in 12.5 µL of Fc blocking buffer, and incubated on ice for 15 minutes. Cells were stained for 20 minutes on ice in 25 µL total volume of antibodies in the Fc blocking buffer. The following antibodies were used at the following dilutions: anti-CD62L, 1:1200 (eBioscience, clone MEL-14, APC-eFluor780), anti-CD44, 1:1200 (eBioscience, clone IM7, PerCP-Cyanine5.5), anti-CD25, 1:1200 (BioLegend, clone PC61, BV510), anti-CD366/Tim-3, 1:100 (BioLegend, clone RMT3-23, APC), anti-CD127, 1:100 (eBioscience, clone A7R34, PE). After staining, cells were centrifuged, resuspended in FACS buffer, and analyzed on an Attune NxT system (ThermoFisher). Data were analyzed using FlowJo software (BD). Statistical analysis was performed using R. For all experiments, n=2 or 3 biological replicates. Replicates are indicated as individual dots in plots.

### *In vivo* CD8^+^ T cell effector differentiation study in the context of viral infection

5- to 7-week-old female C57BL/6 mice were purchased from Jackson Laboratory (Bar Harbor, ME, USA) for this study. H2-D^b^GP33-specific Thy1.1^+^ TCR-transgenic P14 mice were maintained in our colony. Mice were used in accordance with Institutional Animal Care and Use Committee (IACUC) guidelines. Naïve ∼2000 GP33-specific P14 CD8^+^ T cells were adoptively transferred into B6 mice, followed by intravenous infection with Lymphocytic choriomeningitis virus (LCMV) (2×10^6^ PFU LCMV_Cl13_). Three PBS, mIL-21, and 21h10, were administered daily intraperitoneally at 2.5 µg per mouse from day 0 to day 6 after infection. Mice were euthanized on day 7 post-infection. Blood and spleen were collected from each mouse, and antigen-specific CD8^+^ T cells were analyzed for effector molecule granzyme B expression using flow cytometry by gating GP33-specific P14 CD8^+^ T cells using the Thy1.1 congenic marker.

### CAR-T cell generation and maintenance

The retroviral vector expressing the CD19 chimeric antigen receptor (CAR) was constructed using sequences from the clone FMC63 human CD19 single chain variable fragment, and the murine CD28 and CD3ζ sequences. A Myc tag was added to the N-terminus of the CAR construct. After the CAR sequence, a T2A protein cleavage site was inserted, followed by Thy1.1 marker sequence. This CAR construct was then cloned into murine retroviral vector MP71. The Plat E retroviral packaging cell line was used to generate retroviruses following transfection with the CAR construct using lipofectamine. Retroviral supernatant was collected 48 hours after transfection, and used for transduction of CD8^+^ T cells in the presence of polybrene using spinfection (2000g for 60 minutes at 37°C). Prior to transduction, CD8^+^ T cells were purified with MojoSort murine CD8^+^ T cell isolation kit (BioLegend Cat# 480007) and activated by plate-bound anti-CD3/CD28 in the presence of 10 ng/mL IL-7 and 10 ng/mL IL-15 for 24 hours. 16 hours after the transduction, the retroviral media was removed and the cells were transferred to uncoated 6-well plates and expanded in media containing 10 ng/mL IL-7 and IL-15 with PBS, mIL-21 (100 ng/ml) or 21h10 (100 ng/ml) for 3 days.

### Serial tumor killing assay by CAR T cells using IncuCyte

MC38 adenocarcinoma tumor line was obtained from ATCC, and transduced using lentivirus to express human CD19 truncated (hCD19t) antigen and eGFP. The line was clonally selected and expanded for use in IncuCyte assay. For the Incucyte assay, MC38-CD19t eGFP cells were resuspended and plated at 25,000 cells per well in a clear flat bottom 96-well plate. 2 hours later, T-cells were added at 45,000 CAR-positive cells per well in triplicates. 25,000 MC38-CD19t eGFP cells tumor cells were added to the wells again at 36- and 68-hour time-points with half-volume media changes containing 21h10, mIL-21, and IL-7+IL-15. Tumor cell lysis was imaged in IncuCyte S3 every 2 hours and quantified by the loss of GFP signal (Mean Green Image). Acquisition time was set to 300 milliseconds, and Mean Green Image was used to evaluate cytotoxicity of all treatments.

### Cancer cell lines and cell culture

Wild-type B16F10 melanoma cells (ATCC) were cultured in DMEM supplemented with 10% fetal bovine serum (Life Technologies Cat# 26140079), 1% GlutaMAX (Gibco Cat# 35050061), and 1% penicillin-streptomycin (Gibco Cat# 15140122). MC38 murine colon adenocarcinoma cells from James W. Hodge and Jeffrey Schlom’s labs were cultured in DMEM supplemented with 10% fetal bovine serum (Life Technologies Cat# 26140079), 1% penicillin-streptomycin, 1% MEM NEAA (Gibco Cat# 11140050), 1% Sodium Pyruvate (Gibco Cat# 11360070), and 1% HEPES (Gibco Cat# 15630080). All cells were trypsinized and split every 3 days to avoid over-confluency and were maintained at 37°C in a humidified incubator with 5% CO_2_. Cells used for *in vivo* experiments had been passaged for less than 2 months, negative for known murine pathogens, and were implanted at >95% viability.

### Syngeneic murine tumor model experiments

All animal protocols were approved by the Dana-Farber Cancer Institute Committee on Animal Care (Protocol 14-019 and 14-037) and are in compliance with the NIH/NCI ethical guidelines for tumor-bearing animals. At day 0, 6-8 week-old C57BL/6J mice (JAX Cat# 000664) were inoculated with 80-90% confluent tumor cells (cell line is indicated per experiment). Starting on day 3-5, mice were treated daily with the listed test items, and 21h10 was injected daily following each regimen indicated per experiment. Mice were monitored for survival, weight change, and symptoms of toxicity, including pallor, noticeable weight loss, and fatigue. Mice were euthanized if they lost 20% of body weight or their tumors ulcerated or reached 2,000 mm^3^ in volume, which is the maximal permitted tumor size for these studies.

### *In vivo* MC38 adenocarcinoma cancer experiments

For the antitumor efficacy experiment in MC38 adenocarcinoma cancer, C57BL/6 mice were inoculated subcutaneously with 500,000 MC38 adenocarcinoma cells. 5 days post-tumor inoculation, mice were dosed intraperitoneally with 21h10 (15 µg per mouse, n=5 mice), 21AT36 (13.9 µg per mouse, n=5 mice), anti-PD-1 (Bioxcell Cat# BE0146; 150 µg per mouse, n=5 mice), diluted in sterile endotoxin-free PBS, daily for two weeks. All mice were monitored for overall tumor growth and survival.

For the comparison experiment with IL-21, C57BL/6 mice were inoculated subcutaneously with 500,000 MC38 adenocarcinoma cells. When tumors were palpable 6 days post-tumor inoculation, mice were dosed intraperitoneally with PBS (n=5 mice), 21h10 (15 µg per mouse, n=5 mice ), 1x mIL-21 (14.9 µg per mouse, n=5 mice; R&D Systems Cat# 594-ML-100/CF), 10x mIL-21 (149 µg per mouse, n=5 mice), diluted in sterile endotoxin-free PBS, daily for 5 days. Tumors were measured every other day to monitor tumor growth.

For the rechallenge experiment, mice that had previously cleared their MC38 tumors after treatment with 21h10 (15 µg per mouse, n=56 mice) or anti-PD-1 (150 µg per mouse, n=8 mice) and were tumor-free for 30-90 days were used. Two days before and on the day of tumor inoculations, mice were treated with anti-CD4 (150 µg per mouse; Bioxcell Cat# BE0003-1) and/or anti-CD8 depleting antibodies (150 µg per mouse; Bioxcell (valid) # BE0061) or rat IgG2b isotype (150 µg per mouse; clone LTF-2, BioXCell Cat# BE0090) diluted in sterile endotoxin-free PBS. Then, the tumor-free mice and naive mice were inoculated subcutaneously with new MC38 tumors, 1 million cells per mouse. Tumors were measured every other day to monitor tumor growth. Tumor measurements above 200 mm^3^ were considered the endpoint.

To examine the role of anti-21h10 antibodies in anti-tumor activity, wild-type mice were treated daily with 21h10 (15 µg per mouse, n=38 mice) for 9 days and then taken off treatment for 9 days for 4 cycles, allowing mice to produce high levels of anti-21h10 antibodies. At the end of each treatment cycle, blood serum was collected from mice and used to measure anti-21h10 antibody levels via an ELISA (see methods outlined below). On the ELISA readings, samples were categorized as having low titer antibodies if their absorbance values were higher than two standard deviations above the mean values of the PBS samples; samples were categorized as having high titer antibodies if their absorbance values were higher than two times this value. Mice with varying anti-21h10 antibodies were inoculated subcutaneously with 500,000 MC38 adenocarcinoma cells. When tumors were palpable 6 days post-tumor inoculation, mice were dosed intraperitoneally with PBS (n=6 mice with anti-21h10 antibodies, n=5 naive mice) or 21h10 (15 µg per mouse, n=32 mice with anti-21h10 antibodies, n=15 naive mice) daily for 17 days. Tumors were measured every other day to monitor tumor growth.

To demonstrate the dependence of 21h10 on IL-21 signaling, wild-type and *Il21r*-/- mice (Jackson labs stock #019115) were inoculated subcutaneously with 1,000,000 MC38 adenocarcinoma cells. 6 days post-tumor inoculation, mice were dosed intraperitoneally with 21h10 (15 µg per mouse, n=5 wild type mice and n=4 *Il21r*-/- mice), 21AT36 (13.9 µg per mouse, n=5 wild type mice and n=4 *Il21r*-/- mice) , anti-PD-1 (150 µg per mouse, n=4 wild type mice and n=3 *Il21r*-/- mice), diluted in sterile endotoxin-free PBS, daily for two weeks. All mice were monitored for overall tumor growth and survival.

To examine the role of B cells in the anti-tumor activity of 21h10, wild type and μMT mice were inoculated subcutaneously with 1,000,000 MC38 adenocarcinoma cells. 5 days post tumor-inoculation, mice were dosed intraperitoneally with 21h10 (15 µg per mouse, n=5 wild type mice and n=5 μMT mice) or PBS daily for two weeks. All mice were monitored for overall tumor growth and survival.

### *In vivo* adoptive cell transfer experiments

For comparison experiments with mIL-21, mice received CD8^+^ T cells from TRP1^high^ (Jackson labs stock #030958) and TRP1^low^ (Jackson labs stock #030957) sex-matched donor mice via intravenous injection on day -1. The mice were inoculated subcutaneously with 500,000 wild-type B16F10 cells on day 0. Starting on day 5, mice were injected daily intraperitoneally with PBS (n=16 mice) or therapeutic doses of mIL-21 (14.9 µg per mouse; n=11 mice), 21h10 (15 µg per mouse, n=11 mice) diluted in sterile endotoxin-free PBS. Final treatments were given on day 25, and mice were monitored for overall tumor growth and survival. Mice with tumors that ulcerated before reaching a size of 500mm^3^ were excluded from the dataset.

For the comparison experiment with Neo-2/15, half of the mice received CD8^+^ T cells from TRP1^high^ and TRP1^low^ sex-matched donor mice via intravenous injection on day -1. All mice were inoculated subcutaneously with 500,000 wild-type B16F10 cells on day 0. Starting on day 5, all mice were injected daily with PBS (n=10 for each group with and without adoptive transfer) or therapeutic doses of Neo-2/15 (30 µg per mouse, n=10 for each group with and without adoptive T cell transfer) or 21h10 (15 µg per mouse, n=15 for group without adoptive transfer, n=13 for group with adoptive T cell transfer) diluted in sterile endotoxin-free PBS. Final treatments were given on day 21 and day 18 for mice that did and did not receive adoptive transfer of TRP1 cells, respectively. All mice were monitored for overall tumor growth and survival. Mice with tumors that ulcerated before reaching a size of 500mm^3^ were excluded from the dataset.

For the comparison experiment with TNFα blockade, mice received CD8^+^ T cells from TRP1^high^ and TRP1^low^ sex-matched donor mice via intravenous injection on day -1. The mice were inoculated subcutaneously with 500,000 wild-type B16F10 cells on day 0. Starting on day 5, mice were treated intraperitoneally with PBS (n=10 mice ) or 21h10 (15 µg per mouse, n= 20 mice) daily and rat IgG2b isotype (150 µg per mouse,) or anti-TNFα (150 µg per mouse) every 3 days for 21 days. Tumors were measured every other day to monitor tumor growth.

### *In vivo* tumor-infiltrating T cell analysis of Foxp3-GFP mice

On day 0, Foxp3-GFP mice (JAX Cat# 006772) received T cells from TRP1^high^ and TRP1^low^ CD45.1 donor mice and were inoculated subcutaneously with 500,000 wild-type B16F10 cells. Starting on day 5 after tumor cell inoculation, mice were injected daily (intraperitoneal) with therapeutic doses of Neo-2/15 (30 µg per mouse), 21h10 (15 µg per mouse), and mIL-21 (R&D Systems Cat# 594-ML) (14.9 µg per mouse). On day 14 post-inoculation, mice were treated with Brefeldin A (Sigma Aldrich Cat#B7651) intraperitoneally. 6 hours later, tumors were collected and homogenized into single-cell suspensions (and spleen briefly suspended in ACK lysis buffer) and cell populations were analyzed by flow cytometry using the following antibodies: Brilliant Violet 510 anti-CD4 (clone RM4-5, BioLegend Cat# 100559), Brilliant Violet 785 anti-CD8a (clone 53-6.7, BioLegend Cat# 100750), PE anti-CD45.1 (clone A20, BioLegend Cat# 110708), Brilliant Violet 711 anti-CD45.2 (clone 104, BioLegend Cat# 109847), Pacific Blue anti-CD11b (clone M1/70, BioLegend Cat# 101224), PE/Cyanine7 anti-PD-1 (clone RMP1-30, BioLegend Cat# 109109), and APC anti-TIM3 (clone RMT3-23, BioLegend Cat# 119706). All antibodies were used at a 1:100 dilution. Populations of adoptively transferred cells were distinguished by the congenic CD45.1 marker from the host’s endogenous CD45.2^+^ cells.

### Intracellular cytokine staining of tumor-bearing mice

Additionally, tumor samples from above were analyzed using the following extracellular antibodies: Brilliant Violet 510 anti-CD4 (clone RM4-5, BioLegend Cat# 100559), Brilliant Violet 785 anti-CD8a (clone 53-6.7, BioLegend Cat# 100750), PE anti-CD45.1 (clone A20, BioLegend Cat# 110708), Brilliant Violet 711 anti-CD45.2 (clone 104, BioLegend Cat# 109847), Pacific Blue anti-CD11b (clone M1/70, BioLegend Cat# 101224), and also were processed with Intracellular Staining Permeabilization Wash Buffer (BioLegend Cat# 421002) and Fixation Buffer (BioLegend Cat# 420801) according to the manufacturer’s instructions and stained with intracellular antibodies: Brilliant Violet 421 anti-IFN-𝛾 (clone XMG1.2, BioLegend Cat# 505830), and Alexa Fluor anti-granzyme B (clone GB11, BioLegend Cat# 515405). Populations of adoptively transferred cells were distinguished by the congenic CD45.1 marker from the host’s endogenous CD45.2^+^ cells.

### ELISA serum analysis for 21h10 specific anti-drug antibodies

Serum from mice treated daily with PBS, 21h10, and mIL-21 were tested for formation of 21h10 specific anti-drug antibodies. Mice were bled at the endpoint for each experiment. Individual whole blood samples were incubated for 5-10 minutes at 25°C, spun at 5,000 rpm for 5 minutes at 25°C, and the serum layer was collected. For the subsequent ELISAs, 96-well high-binding plates were coated overnight at 4°C with 21h10 protein (500 ng per well) diluted in 1X coating buffer (from ELISA set, Biolegend Cat#430804), and blocked with PBS + 10% IFS for 24 hours at 4°C. Serum samples diluted 1:100 in 1X Assay Diluent A (from ELISA set, Biolegend Cat# 430804) were plated and incubated for 2 hours at 25°C. Binding was detected with anti-mouse IgG HRP-linked antibody (Cell Signaling Cat# 7076) and incubated at 25°C for 1 hour and developed with 3,3′,5,5′-Tetramethylbenzidine (TMB) (Sigma Aldrich Cat# T0440). The reaction was stopped with 1N HCl (Fisher Scientific Cat# SA48500) after 5-10 minutes, and intensity measurement was read at 450 nm. Serum samples positive for anti-drug antibodies (above the dotted line) were determined to have an intensity measurement 2 SD above the mean absorbance value of samples from mice treated with PBS. Fractions indicate the number of mice classified as positive for anti-drug antibodies out of the total mice in each treatment group.

### *In vivo* anti-TRP1 TA99 antibody experiment

For melanoma experiment with 21h10 and anti-TRP1 TA99 antibody, C57BL/6 mice were inoculated subcutaneously in the flank with 80-90% confluent wild-type B16F10 cells (500,000 cells per mouse) on day 0. Starting on day 3, mice were administered bi-weekly treatment with TA99 antibody intraperitoneally (150 µg per mouse). Starting on day 5, mice were injected daily with therapeutic doses of 21h10 (15 µg per mouse).

### *Ex vivo* adoptive cell transfer

For *ex vivo* priming experiment with 21h10 and Neo-2/15, C57BL/6 mice were inoculated subcutaneously in the flank with 80-90% confluent wild-type B16F10 cells (500,000 cells per mouse) on day 0. On day 0, CD8^+^ T cells were harvested from a TRP1^high^ donor mouse and plated together with 21h10 (500 nM) or Neo-2/15 (500 nM) in RPMI complete medium in a 6-well plate that was pre-coated overnight at 4°C with anti-CD3 antibody (BioLegend Cat# 100340) at 100 pg/mL. After 2 days, cells were harvested and transferred by intravenous injection into the tumor-bearing mice. 10 mice were included in the PBS control group, 5 in the Neo-2/15-treated group, and 10 in the 21h10-treated group.

### *In vivo* toxicity experiments

For testing treatment toxicity *in vivo*, C57BL/6, *Rag2-/-*, or *Rag2-/- Il2rg-/-* mice were treated with PBS or 21h10 (15 µg per mouse) daily and with rat IgG2b isotype (150 µg per mouse, clone LTF-2, BioXCell Cat# BE0090) or IgG2a anti-mouse NK1.1 depleting antibodies (150 µg per mouse, clone PK136, BioXCell Cat# BE0036) every three days. Treatment was continued for each mouse until weight loss reached 20% of the initial starting weight and mice were sacrificed. Serum from these mice was collected and analyzed by cytokine bead array (Evetechnologies). Cytokine values from WT and *Ifng*-/- mice were compared to *Rag2*-/- mice with a Wilcoxon rank test followed by P-value adjustment with the Benjamini-Hochberg (BH) procedure.

### Western blot analysis for the *in vivo* stability of 21h10

C57BL/6 mice were treated with 21h10 (n=12 mice), mIL21 (n=12 mice), or PBS (n=4 mice) via intravenous tail vein injections. 1, 6, 9, and 24 hours following treatment, mice were bled and euthanized; spleens and lymph nodes were harvested and lysed in a buffer composed of 50 uM NaCl, 50 uM HEPES, 0.5% NP-40 with protease and phosphatase inhibitors in deionized water. A bicinchoninic acid assay was performed to determine the protein concentration of lysates. Equal amounts of spleen lysates were loaded in 4 to 20% SDS-polyacrylamide gels followed by a transfer to a polyvinylidene difluoride (PVDF) membrane using Bio-Rad Trans-blot Turbo Transfer System. Detection antibodies were incubated in 3% BSA in Tris-buffered saline with Tween 20 (TBS-T) composed of tris buffer, NaCl, and Tween 20. The antibodies used were: Phospho-Stat3 (9145S) and Stat3 (12640S). Once images of western blots were obtained, the relative density and size of bands were quantified on ImageJ.

### Single-cell RNA-sequencing preparation

Mice bearing B16F10 tumors and adoptively transferred TRP1-specific T cells (as above) were sacrificed at 14 days post-tumor inoculation. Tumors were collected from mice as described previously^70^. Briefly, tumors were removed and manually dissociated with scissors and incubated in RPMI with tumor digestion enzymes (Miltenyi Cat# 130-096-730) for 30 minutes at 37°C. Digested tumors were filtered through a 40 μM strainer and the single-cell suspension was resuspended in PBS (with 0.5% BSA and 2 mM EDTA) and incubated with CD45 MicroBeads (Miltenyi Cat# 130-052-301). Following the manufacturer’s protocol, magnetically enriched CD45^+^ populations were collected and counted with a hemocytometer. For each tumor from a treatment group, a different TotalSeqC hashing antibody (mouse Hashtags 1-5, Biolegend) was incubated with 2 μL Ab/million cells in PBS (with 2% FBS) for 20 minutes at 4°C. Following three washes with 0.05% ultrapure BSA (Invitrogen Cat# AM2618) in PBS, cells were counted again and resuspended at 1,000 cells/μL. Approximately 4,000 cells/tumor were pooled across each treatment group (five hashtags/treatment group) and loaded onto a 10X Chromium Controller with a Single Cell K Chip (PN-2000182) using the Chromium Next GEM Single Cell 5ʹ GEM Kit v2 reagents and beads (PN-1000244 and PN-1000264). Library preparation was performed with the Library Construction Kit (PN-1000190) to generate gene expression libraries and 5’ Feature Barcode kit (PN-1000256) was used to generate hashtag libraries. Samples were sequenced on an Illumina NovaSeq 6000 instrument with 2×150bp sequencing (Azenta Life Sciences).

### Single-cell RNA-sequencing pre-processing

The *Mus musculus* genome fasta and General Transfer Format (GTF) annotation files were downloaded from the Ensembl database (release 108). Cellranger (v7.0.0) ‘mkgtf’ was used to filter the GTF file using the defaults from 10x Genomics. A custom reference was generated by first appending the sequences for the TRP1^high/low^ rearranged α/β TCR chains^45^ to the *M. musculus* fasta file, adding TRP1 TCR annotations to the filtered GTF file, and finally using these two new files as inputs for Cellranger ‘mkref’ to build the reference. Cellranger ‘count’ was used to align reads to this new reference (Gene Expression) and hashtag antibody sequences (Antibody Capture). The unfiltered Cellranger count matrices were imported into R using the *Seurat* (v4.3.0) package. The *DropletUtils* (v1.18.1) package ‘emptyDrops’ function was used to exclude potential empty droplets from the count matrix^71^ using a cutoff of FDR < 0.01. Genes expressed in fewer than five cells were excluded from downstream analysis. Hashtag reads were used to demultiplex samples with the Seurat ‘HTODemux’ function, using a kmeans function with “nstarts = 30” and a 99% positive quantile threshold. Only cells classified as a “singlet” but not as a “doublet” or “negative” by hashtag demultiplexing were kept for downstream analysis. Following hash demultiplexing, cells were further filtered out if they contained fewer than 1,000 genes per cell or more than 10% mitochondrial (*mt-*) reads.

### Single-cell RNA-sequencing clustering

Each filtered, demultiplexed sample was log normalized, and 2,000 variable features were identified and used to find 3,000 integration features to combine samples to minimize batch effects^72^. The newly integrated dataset was scaled, and a principal component analysis (PCA) was performed, from which the top 25 PCs were used for uniform manifold approximation and projection (UMAP) generation. From the 25 PCs, a shared nearest neighbor (SNN) graph was constructed (k = 20 neighbors), and clusters were classified by the Louvain algorithm. The ‘FindAllMarkers’ function was used to find the top cluster-defining genes, and these gene lists were used to assign cell types.

### TRP1 identification and T cell sub-clustering

From the custom reference built above, TRP1^high^ α/β and TRP1^low^ α/β TCR chains were counted by Cellranger. The sum of α and β chain expression for TRP1^high^ or TRP1^low^ TCRs was calculated for each cell. Cells were classified as TRP1^high^ or TRP1^low^ if they had an exclusive expression of one clonotype but not the other (e.g., high = TRP1^high^ > 0 & TRP1^low^ = 0). The few cells that had counts associated with both TCRs were not considered to be TRP1^high^ or TRP1^low^. T cell sub-clustering was performed by subsetting cells expressing at least two CD3 chains (*Cd3d/e/g*). This initial T cell subset was split by sample and then new variable features were identified, with the exclusion of TCR variable genes (*Tr[abgd][vj]*), to reintegrate the samples back together. The top 30 PCs were used for UMAP and SNN graph generation. Six clusters separated from the main population, exemplified by above-average expressions of *Csf1r*, *Sirpa*, *H2-DMb2*, and *Lyz2*. These contaminating non-T cell clusters were removed from the initial T cell sub-clustering, and the process of splitting and reintegrating was repeated to yield the final T cell sub-clusters, with the added step of regressing out ribosomal (*Rp[sl]*) and mitochondrial reads when scaling the data. Sub-clusters were identified as above with Louvain community detection and ‘FindAllMarkers.’ CD4^+^ T cell sub-clustering was performed by selecting cells in the “CD4” and “Treg” sub-clusters or cells with the exclusive expression of *Cd4* (*Cd4* ≥ 1 and *Cd8a/b1* = 0). The same workflow of splitting, reintegrating (excluding TCR variable gene expression), and scaling while regressing out ribosomal/mitochondrial reads was performed. CD4^+^ T cell sub-clusters were identified as above.

### Differential gene expression analysis

Across each T cell sub-cluster, we performed a pseudo-bulk method of differential gene expression analysis^73,74^. Each mouse within a treatment group was treated as a replicate, and count matrices were summed across genes for each sub-cluster per mouse. Using the *DESeq2* (v1.38.3) package^75^ and pseudo-bulk gene expression values, differentially expressed genes were identified across treatment groups and sub-clusters. For gene changes in TRP1-specific T cells, cells classified as either TRP1^high^ or TRP1^low^ cells were subset and counts across each gene were summed for each mouse for pseudo-bulk analysis using *DESeq2*. For Gene Set Enrichment Analysis (GSEA), fold changes within the T cell subset object were calculated with the *Seurat* ‘FoldChange’ function, comparing each specified treatment group. Fold changes were used with the *fgsea* (v1.24.0) package along with the *Molecular Signatures* (v7.2) Hallmark gene signatures^76^ for GSEA.

### Organotypic tumor spheroid preparation and microfluidic device culture

PDOTS were generated as previously described^48,77^. Briefly, fresh tumor specimens from patients were received in full DMEM media (Corning, 10-013-CV), with 10% BenchMark™ Fetal Bovine Serum (GeminiBio, 100-106) and 1% penicillin-streptomycin (Thermo Scientific, 15140122) on ice and minced in a 10 cm plate using sterile forceps and scalpels, if samples arrived late, they were stored in tissue storage media (Miltenyi, 130-100-008) and processed the day after. Minced tumors were resuspended in full DMEM and were passed over 100-μm and 40-μm filters sequentially to generate the S1 (>100 μm), S2 (40-100 μm), and S3 (<40 μm) fractions. S2 and S3 were washed off the filter with fresh full media and were rested in ultra-low attachment plates in (Corning, 3471) in the incubator until loading into the device. S1 fraction was washed from the filter using fresh full media supplemented with 100 U/mL collagenase type IV (Life Technologies, 17104019), and 15 mM HEPES (Gibco, 1560-080). S1 with collagenase was incubated for 15-30 minutes at 37°C followed by the addition of an equal volume of media and subsequent filtering. The S2 fraction was pelleted and resuspended in type 1 rat tail collagen (Corning, 354236) at a concentration of 2.5 mg/mL with 10x PBS + phenol red (Sigma-Aldrich, 114537-5g). A pH of 7.0-7.5 was confirmed with PANPEHA Whatman paper (Sigma-Aldrich, 2629990) after titrating the solution with NaOH (Sigma-Aldrich, 1.09138.1000). The spheroid-collagen mixture (10 μL) was added into the center channel of the AIM 3D microfluidic device (DAX-01, AIM Biotech). Collagen hydrogels with PDOTS were incubated for 20 minutes at 37°C in sterile humidity chambers before being hydrated with media in the side channels with or without treatments. PDOTS were treated with ICB (250 μg/mL pembrolizumab or 240 μg/mL nivolumab + 80 μg/mL relatlimab (opdualag)), human recombinant IL-21, or left untreated.

### PDOTS and MDOTS viability assessment

PDOTS staining and viability analysis was done as previously described^48,77^. In brief, staining was done in microfluidic devices by adding acridine orange/ propidium iodide (AO/PI) solution (Nexcelom, CS2-0106), diluted 1:1 in full media with 12 μg/ml Hoechst (Invitrogen, H3570). After incubation with fluorescent dyes (30 minutes, 37°C), images were acquired using a Nikon Eclipse NiE fluorescence microscope, using X4 lens in 3 colors. Images were analyzed using the NIS-Elements AR software package and live/dead cell quantification was obtained by measuring the total cell area of acridine orange for live cells, PI for dead cells and Hoechst for total cells. Live cells were defined as the total area of the acridine orange channel, or the total area of Hoechst (total cells)-PI (dead cells). Percent change and log2FC (L2FC) data were generated using raw fluorescence data (live) for given treatments relative to control conditions.

## Statistical analysis

All statistics were calculated with GraphPad Prism or R. All error bars are S.E.M. Data were considered significant when p ≤ 0.05; *p < 0.05, **p < 0.01, ***p < 0.001, ****p < 0.0001 (or p-values displayed).

## Data Availability

RNA sequencing has been deposited in the Gene Expression Omnibus (GEO) under the accession number GSE256289. Single-cell RNA sequencing has been deposited in the Gene Expression Omnibus (GEO) under the accession number GSE240834.

## Appendices

### Appendix A. Parameters for scaffold generation

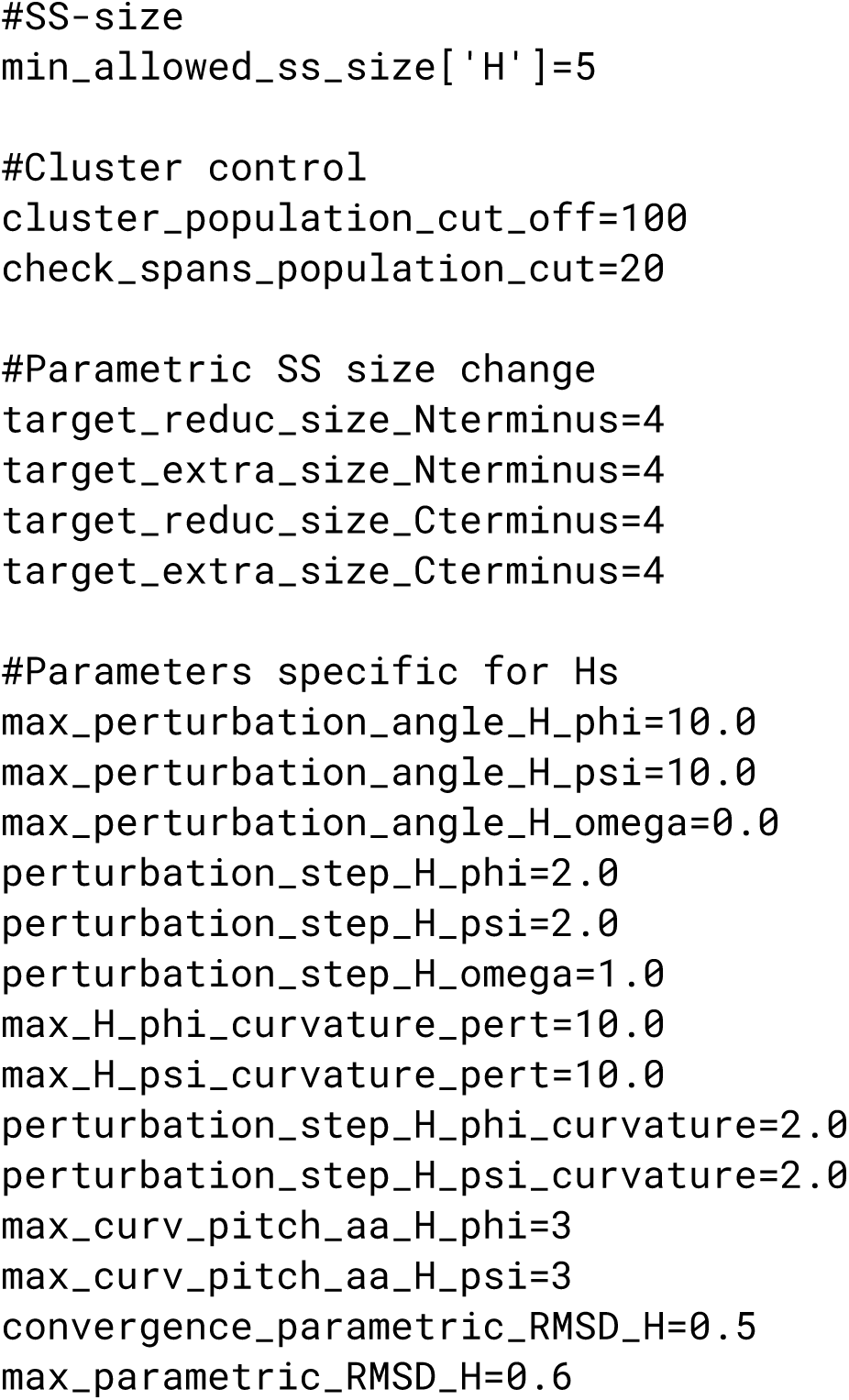

### Appendix B. PDBInfoLabel for scaffold generation and interface design

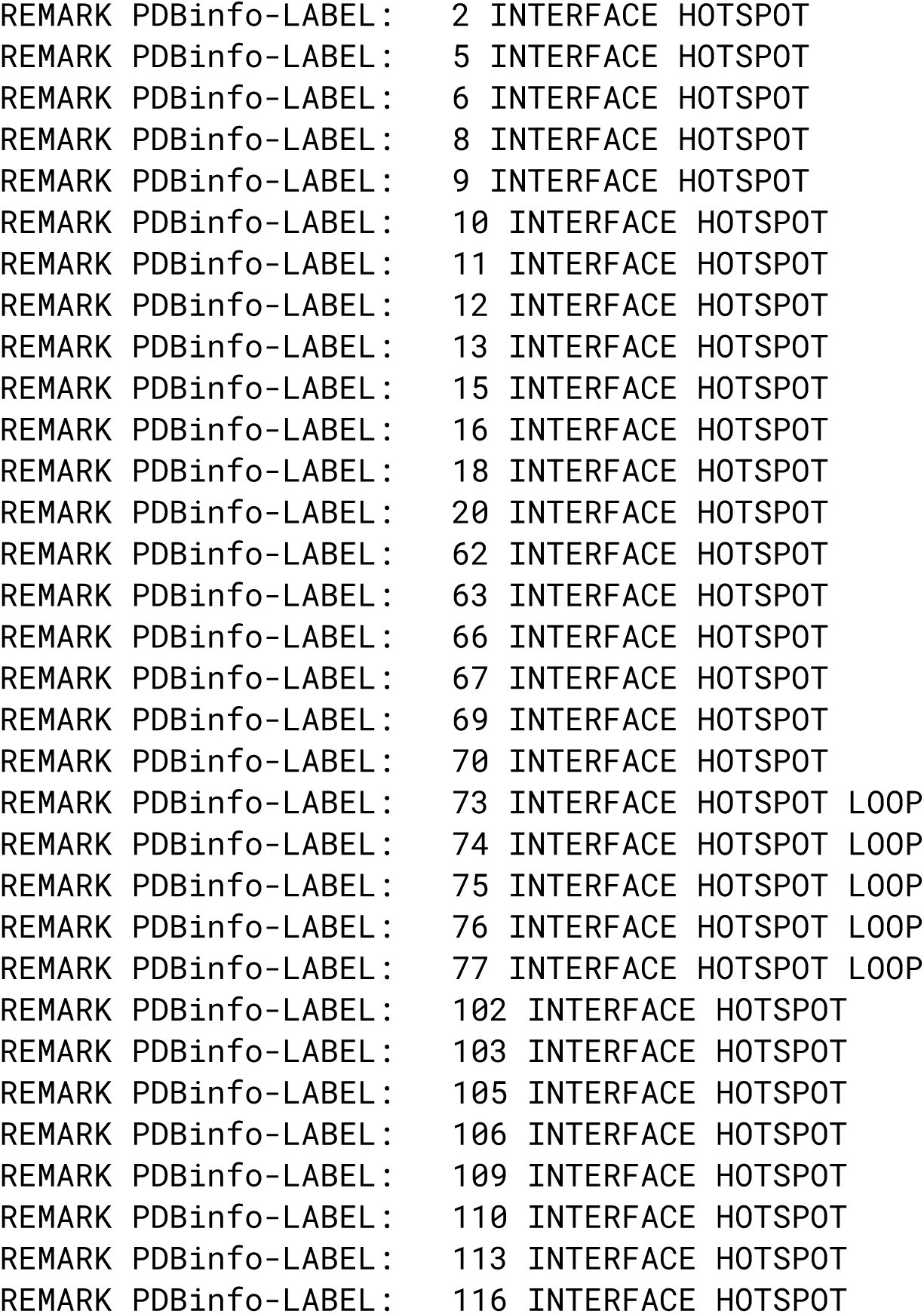

## Acknowledgements

This work was supported with funds provided by The National Cancer Institute (R01CA240339) to J.-H.C., A.Q.R., and D.B.; The National Institute of Allergy and Infectious Disease (R01AI160052) to J.-H.C., S.G., and D.B.; The Division of Intramural Research, the National Heart, Lung, and Blood Institute, the National Institute of Health to S.R., P.L., and W.J.L.; Stanford University Medical Scientist Training Program Grant (T32GM007365) to G.C.A.; Hertz Fellowship to G.C.A.; The National Institute of Health (R01CA177684) to G.C.A. and M.D.; The National Science Foundation Graduate Research Fellowship to E.C.C.; The Institute for Stem Cell & Regenerative Medicine Graduate Fellowship to E.C.C.; The Washington Research Foundation and Translational Research Fund to A.Q.R, and D.S.; The Open Philanthropy Project Improving Protein Design Fund to S.G., M.DeW., L.C., and D.B.; The Audacious Project at the Institute for Protein Design to S.G., M.DeW., P.H., L.C., N.P.K., and D.B.; The National Institute of Biomedical Imaging and Bioengineering, the National Institute of Health, Trailblazer R21 Award (R21EB027327) to H.Y.K.; The National Institute of Health (R21AI154363, R56AG069194, U01CA281848) to V.K.; Hopes and Smiles for Kids Foundation to V.K.; The National Institute of Health (R01AI132819, 5P30CA015704 subaward) to S.S.; Friedman-Rieger Foundation to S.S.; The National Institute of Health (R01AI51321) to K.C.G.; Ludwig Institute for Cancer Research to K.C.G.; Howard Hughes Medical Institute to K.C.G. and D.B.; The National Institute of Health (R01AI169188-01) to S.K.D. and M.D.; The National Institute of Health (R01AI158488-01) to S.K.D.; The Melanoma Research Alliance to S.K.D.; The Peter and Ann Lambertus Family Foundation to M.D.; Additional support provided by the Termeer Early Career Fellowship in Systems Pharmacology to R.W.J.; Generous gift from Robert and Marie McInnes to R.W.J.; The MGH ECOR Fund for Medical Discovery Research Fellowship Award to O.-Y.R.; Funding provided by Massachusetts Life Sciences Center Research Infrastructure Program in support of the Mass General Cancer Center Tumor Cartography Center to R.W.J..

## Author Contributions

J.-H.C. and D.-A.S. performed protein design; J.-H.C. screened, characterized, and optimized the hits; J.-H.C., H.S., and A,Q.-R. measured their binding and stability; J.-H.C., S.G., A.M., P.H., and M.DeW. performed protein expression; J.-H.C., T.N., and L.C. performed BLI for serum stability assay; J.-H.C. performed antagonist design; J.-H.C. and C.J.K. screened and characterized antagonist hits; D.B. supervised protein design, optimization, and *in vitro* characterization; G.C.A. and K.M.J. performed crystallography experiments; K.C.G. supervised crystallography experiments; S.R. performed cell signaling and *in vitro* culture experiments with flow cytometry analysis; S.R. and P.L. performed RNAseq experiments and analysis; W.J.L. supervised cell signaling and RNAseq experiments; E.C.C. performed T cell differentiation experiments; H.Y.K. supervised T cell differentiation experiments; S.T. performed *in vitro* CAR-T cell experiments; A.K. performed LCMV infection experiments and Seahorse analysis of CAR T cells; V.K.. and S.S. supervised CAR-T cell and LCMV infection experiments; B.S.L. and K.Z. performed B16F10 survival experiments; B.S.L., T.A., F.P., and J.G.T., performed MC38 survival and rechallenge experiments; B.S.L., K.Z., and T.A. performed toxicity experiments; T.A. performed anti-drug antibody experiments; B.S.L., T.A., F.P., and J.G.T. performed *in vivo* western blot experiments; B.S.L., M.J.W., K.Z., R.K., and S.Y.L. performed *in vivo* flow cytometry experiments; B.S.L., M.J.W., K.Z., R.K., and S.Y.L. performed scRNA-seq experiments; M.J.W. performed scRNA-seq analysis; B.S.L. performed *in vitro* T cell proliferation experiments; M.D. and S.K.D. supervised *in vivo* cancer experiments; G.M.B. removed the melanoma tissue from patient and established the research protocol; T.S., A.L., and S.C. collected and processed patient melanoma tissue;C.A.P. performed patient melanoma tissue culture; O.-Y.R. designed and ran the PDOTS experiment; R.W.J. supervised the PDOTS experiment and interpreted the data; J.-H.C., B.S.L., S.R., M.J.W., G.C.A., T.A., E.C.C., N.P.K., H.Y.K., V.K., S.S., R.W.J, G.C.A., K.C.G.,

W.J.L., M.D., S.K.D., and D.B. wrote the manuscript.

## Author Information

Graduate Program in Biological Physics, Structure, and Design, University of Washington, Seattle, WA

Jung-Ho Chun

Department of Biochemistry, University of Washington School of Medicine, Seattle, WA

Jung-Ho Chun, Chan Johng Kim, Hojeong Shin, Neil P. King, David Baker

Institute for Protein Design, University of Washington, Seattle, WA

Jung-Ho Chun, Tina Nguyen, Chan Johng Kim, Hojeong Shin, Alfredo Quijano-Rubio, Stacey Gerben, Analisa Murray, Piper Heine, Michelle DeWitt, Umut Y. Ulge, Lauren Carter, Neil P. King, Daniel-Adriano Silva, David Baker

Howard Hughes Medical Institute, University of Washington, Seattle, WA

David Baker

Laboratory of Molecular Immunology and the Immunology Center, National Heart, Lung, and Blood Institute, National Institutes of Health, Bethesda, MD

Suyasha Roy, Peng Li, Warren J. Leonard

Department of Bioengineering and Institute for Stem Cell and Regenerative Medicine, University of Washington, Seattle, WA

Elisa C. Clark, Hao Yuan Kueh

Department of Molecular and Cellular Physiology, Stanford University School of Medicine, Stanford, CA

Gita C. Abhiraman, Kevin M. Jude, K. Christopher Garcia

Howard Hughes Medical Institute, Stanford University School of Medicine, Stanford, CA

K. Christopher Garcia

Ben Towne Center for Childhood Cancer Research, Seattle Children’s Research Institute, Seattle, WA

Asheema Khanna, Samantha Tower, Vandana Kalia, Surojit Sarkar Department of Pediatrics, University of Washington, Seattle, WA Vandana Kalia, Surojit Sarkar

Department of Pathology, University of Washington, Seattle, WA

Surojit Sarkar

Department of Medicine, Division of Gastroenterology, Massachusetts General Hospital, Boston, MA

Birkley S. Lim, Michael J. Walsh, Kevin Zhangxu, Tavus Atajanova, Frank Peprah, Julissa G. Tello, Michael Dougan

Department of Cancer Immunology and Virology, Dana-Farber Cancer Institute, Boston, MA

Birkley S. Lim, Michael J. Walsh, Kevin Zhangxu, Tavus Atajanova, Frank Peprah, Julissa G. Tello, Samantha Y. Liu

Department of Medicine, Harvard Medical School, Boston, MA

Michael J. Walsh, Michael Dougan

Department of Immunology, Harvard Medical School, Boston, MA

Rakeeb Kureshi, Megan T. Hoffman, Stephanie K. Dougan

Mass General Cancer Center, Krantz Family Center for Cancer Research, Department of Medicine, Massachusetts General Hospital, Harvard Medical School, Boston, MA

Or-Yam Revach, Claire A. Palin, Russell W. Jenkins Broad Institute of MIT and Harvard, Cambridge, MA Or-Yam Revach, Genevieve M. Boland, Russell W. Jenkins

Massachusetts General Hospital Cancer Center, Harvard Medical School, Boston, MA

Tatyana Sharova, Aleigha Lawless, Sonia Cohen, Genevieve M. Boland Department of Surgery, Massachusetts General Hospital, Boston, MA Tatyana Sharova, Aleigha Lawless, Sonia Cohen, Genevieve M. Boland

## Competing Interests

J.-H.C., A.Q.-R., U.Y.U., D.-A.S., N.P.K., M.D., S.K.D., and D.B. are co-founders and stockholders of Axxis Bio, Inc., a company that aims to develop the inventions described in this manuscript. J.-H.C., A.Q.-R., U.Y.U., D.-A.S., N.P.K., and D.B. are co-inventors on a US non-provisional patent application (63/378797) and PCT international application (PCT/US23/76260), which incorporates discoveries described in this manuscript. R.W.J. is a member of the advisory board for and has a financial interest in Xsphera Biosciences, Inc., a company focused on using *ex vivo* profiling technology to deliver functional, precision immune-oncology solutions for patients, providers, and drug development companies. R.W.J. has received honoraria from Incyte (invited speaker), G1 Therapeutics (advisory board), Bioxcel Therapeutics (invited speaker). R.W.J. has ownership interest in U.S. patents US20200399573A9 and US20210363595A1. R.W.J.’s interests were reviewed and are managed by Massachusetts General Hospital and Mass General Brigham in accordance with their conflict-of-interest policies. The remaining authors declare no competing interests.

---

**Figure S1.**
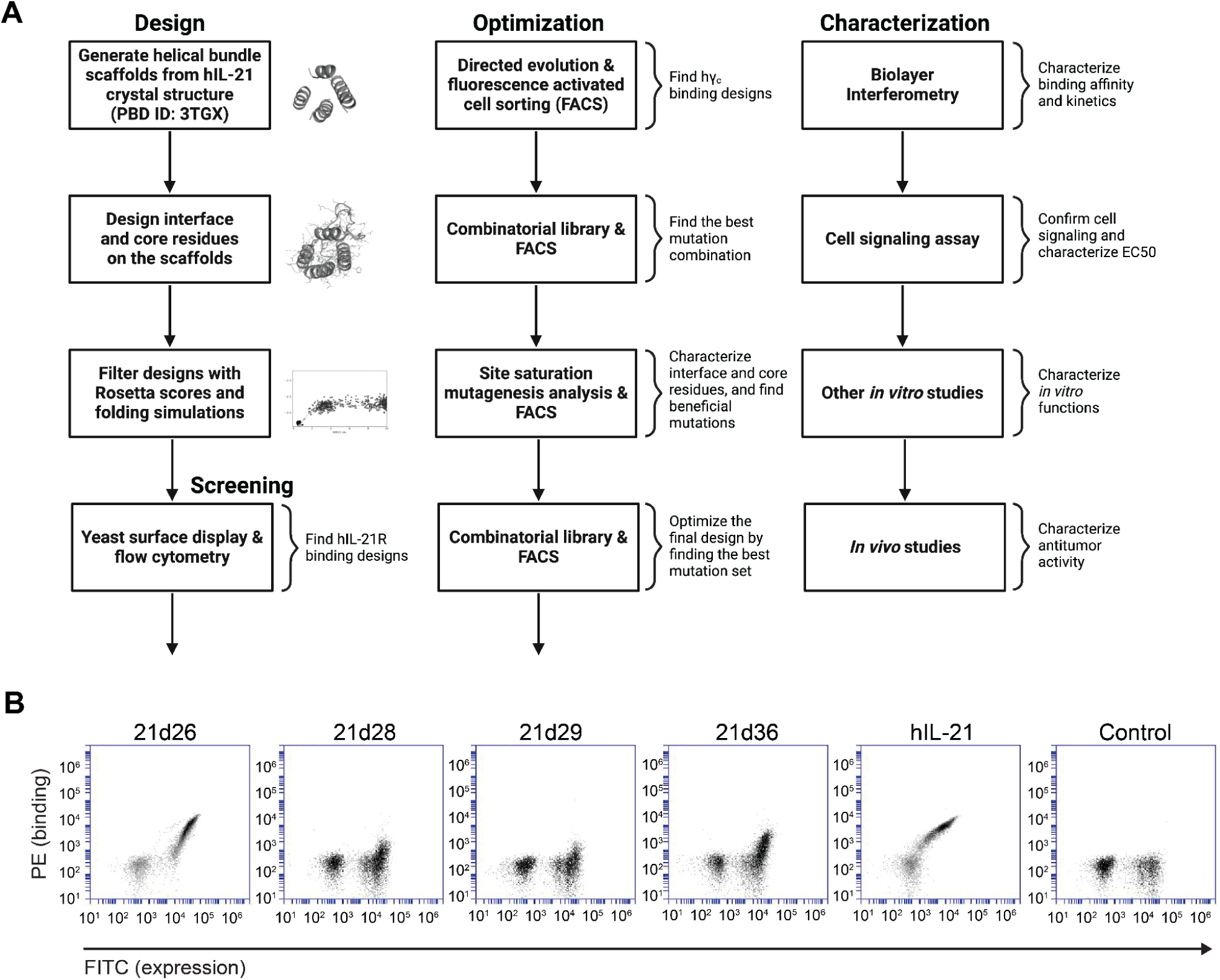
Design, screening, and optimization of IL-21 mimic. (**A**) Stepwise process for designing, screening, optimizing, and characterizing IL-21 mimics. The helical bundle scaffolds are generated from the hIL-21 structure, and their residues are designed in the context of hIL-21R (PDB ID: 3TGX). The designs are computationally screened using *Rosetta* scores and folding simulations. The designs are experimentally screened using yeast surface display with flow cytometry. Optimization processes include directed evolution, SSM analysis, and combinatorial library screening. Further characterizations use BLI, cell signaling assay, and other *in vitro* and *in vivo* assays. (**B**) Four hits (21d26, 21d28, 21d29, and 21d36) from the initial designs that bind to 100 nM of hIL-21R. hIL-21 as positive control.

**Figure S2.**
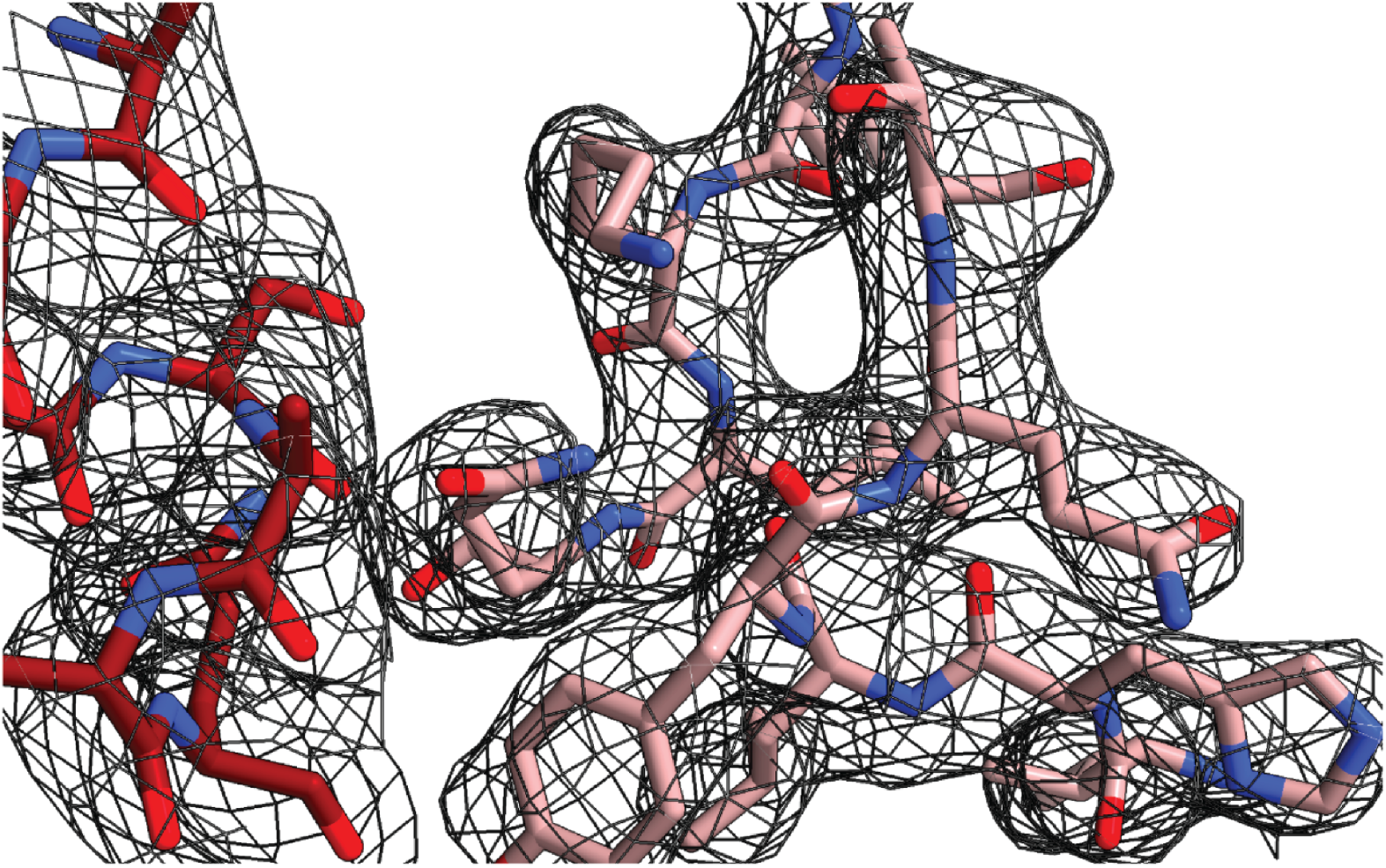
Electron density of 21h10. Representative electron density. 2mFo-Dfc map (shown in gray) contoured at 1.0 σ. 21h10 is shown in red, h𝛾_c_ shown in light pink.

**Figure S3.**
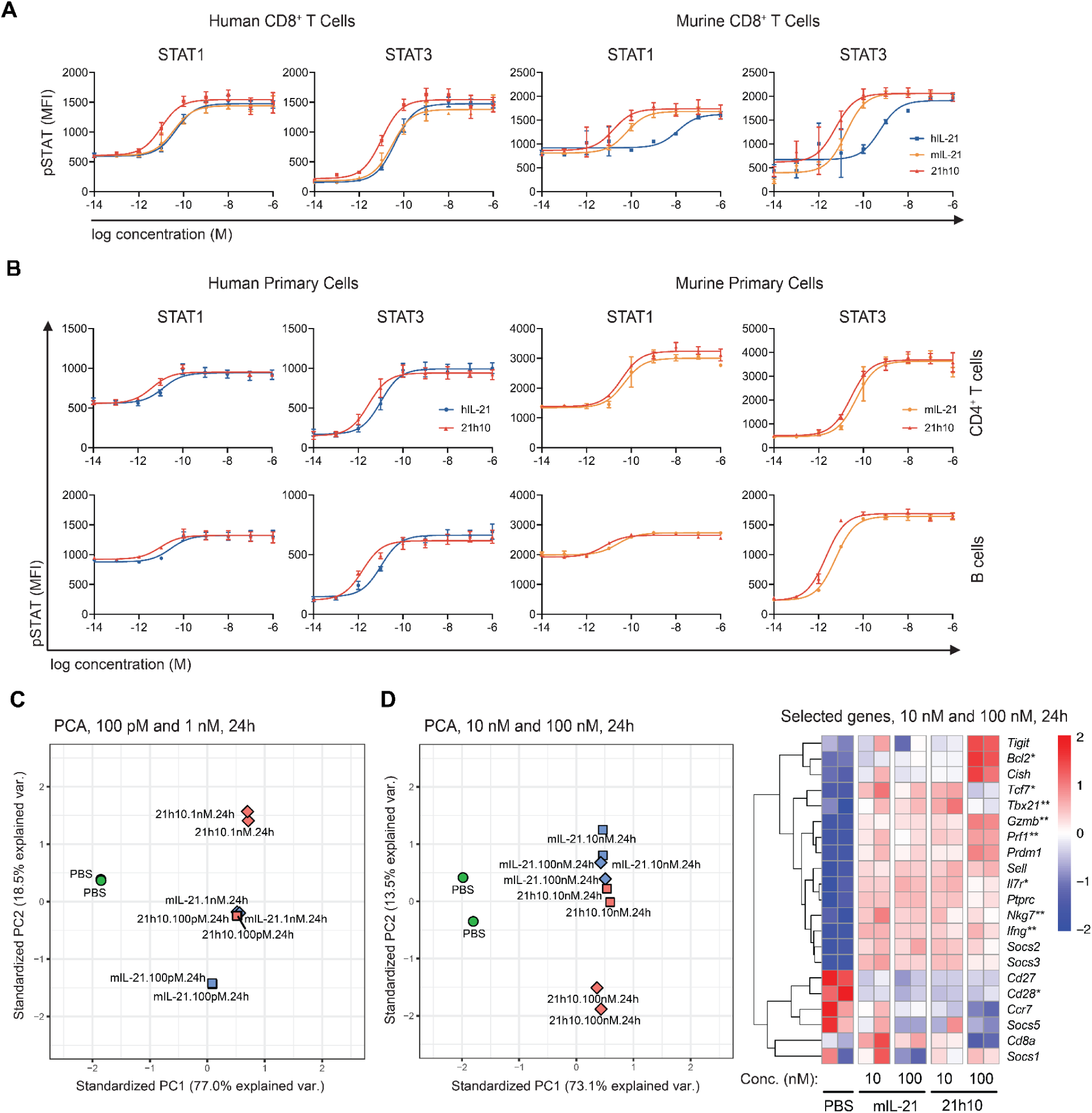
STAT signaling of CD4^+^ and B cells and PCA analyses and gene expression comparison at higher concentrations of cytokines. (**A**) Cross-reactivity of hIL-21, mIL-21, and 21h10 in human and murine CD8^+^ T cells. (**B**) Phospho-STAT signaling of human and murine CD4^+^ T cells and B cells. (**C**) PCA analysis of RNA-seq data from murine CD8^+^ T cells treated with PBS, mIL-21, or 21h10 at 100 pM or 1 nM from Fig. 2B. (**D**) PCA analysis of RNA-seq data of murine CD8^+^ T cells treated with PBS, mIL-21, or 21h10 at 10 nM or 100 nM. A set of genes are compared for expression levels between the treatment groups. * indicates memory-related genes, and ** indicates effector-related genes.

**Figure S4.**
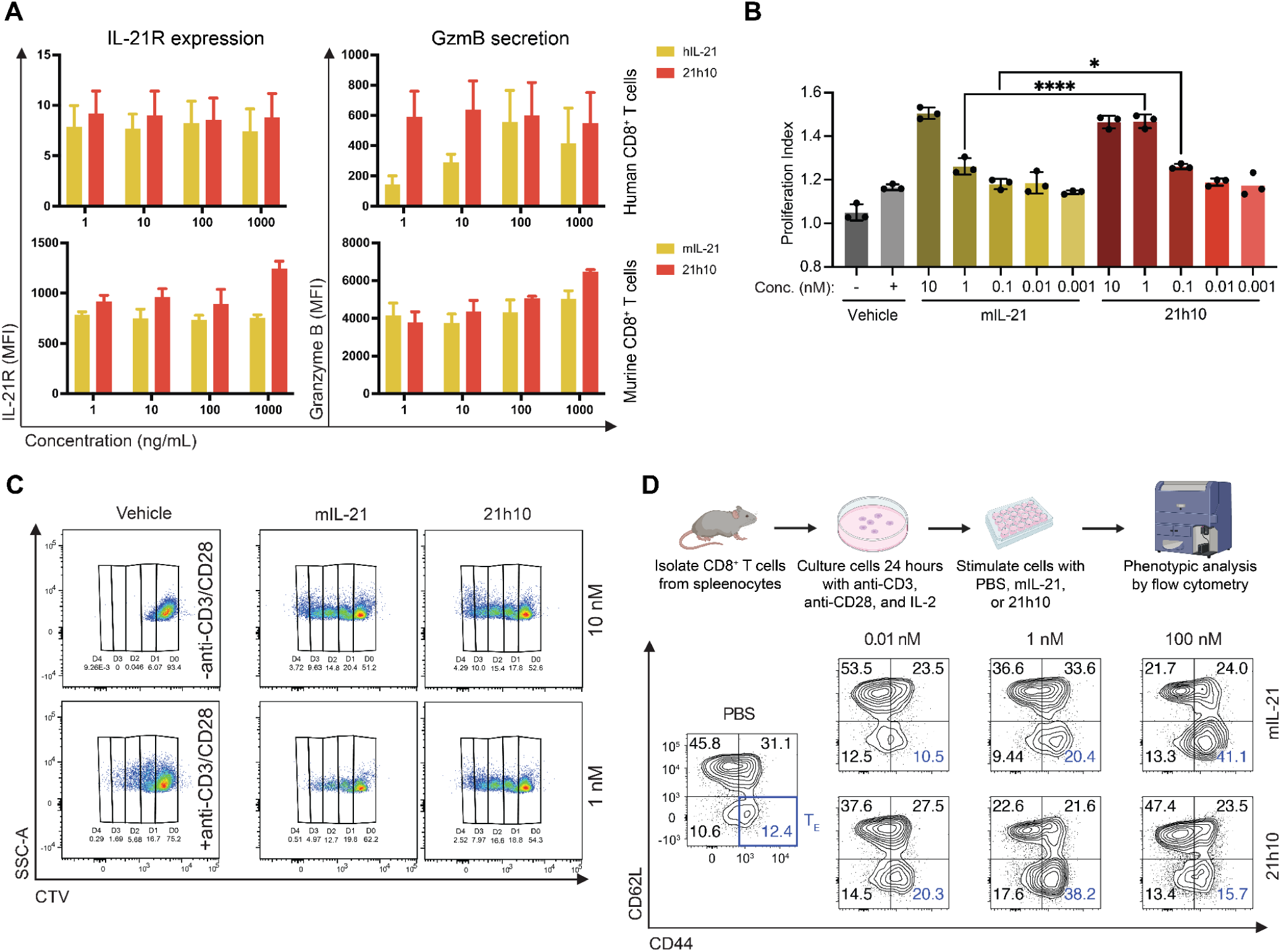
Proliferation, differentiation, and expression of IL-21R and granzyme B of CD8^+^ T cells in the presence of mIL-21 or 21h10. (**A**) IL-21R and granzyme B expressions of murine and human CD8^+^ T cells were measured by flow cytometry after culturing with anti-CD3, anti-CD28, and corresponding cytokines for 72 hours. (**B**) Murine TRP1^high^ CD8^+^ T cell proliferation was quantified using flow cytometry with Cell Trace Violet (CTV). All groups were treated for 72 hours with anti-CD3/CD28 beads along with various amounts of the cytokines that were incubated prior to addition onto the cells at 37°C for 24 hours in human serum. (**C**) Flow cytometry plots of CTV-labeled cells from (**B**). (**D**) Murine CD8^+^ T cells were isolated and then activated with anti-CD3/CD28 antibodies and IL-2 for 24 hours. Cells were then incubated with PBS, mIL-21, or 21h10 at the indicated concentrations for 48 hours. The cells were analyzed for their differentiation into the effector T cell (CD44^+^CD62L^-^) population using flow cytometry.

**Figure S5.**
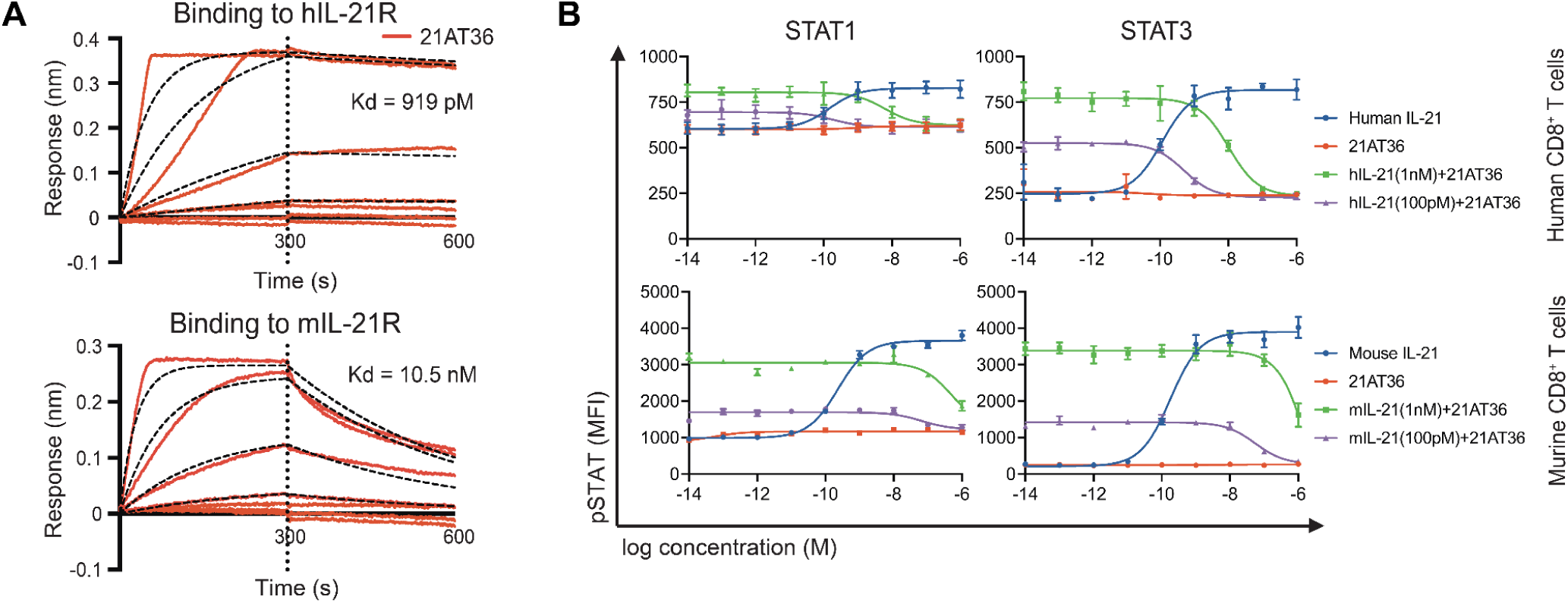
Antagonist, 21AT36, binds to IL-21R but does not signal through STAT. (**A**) Biolayer interferometry against hIL-21R and mIL-21R. (**B**) 21AT36 does not phosphorylate STAT and can compete native IL-21 via IL-21R-binding.

**Figure S6.**
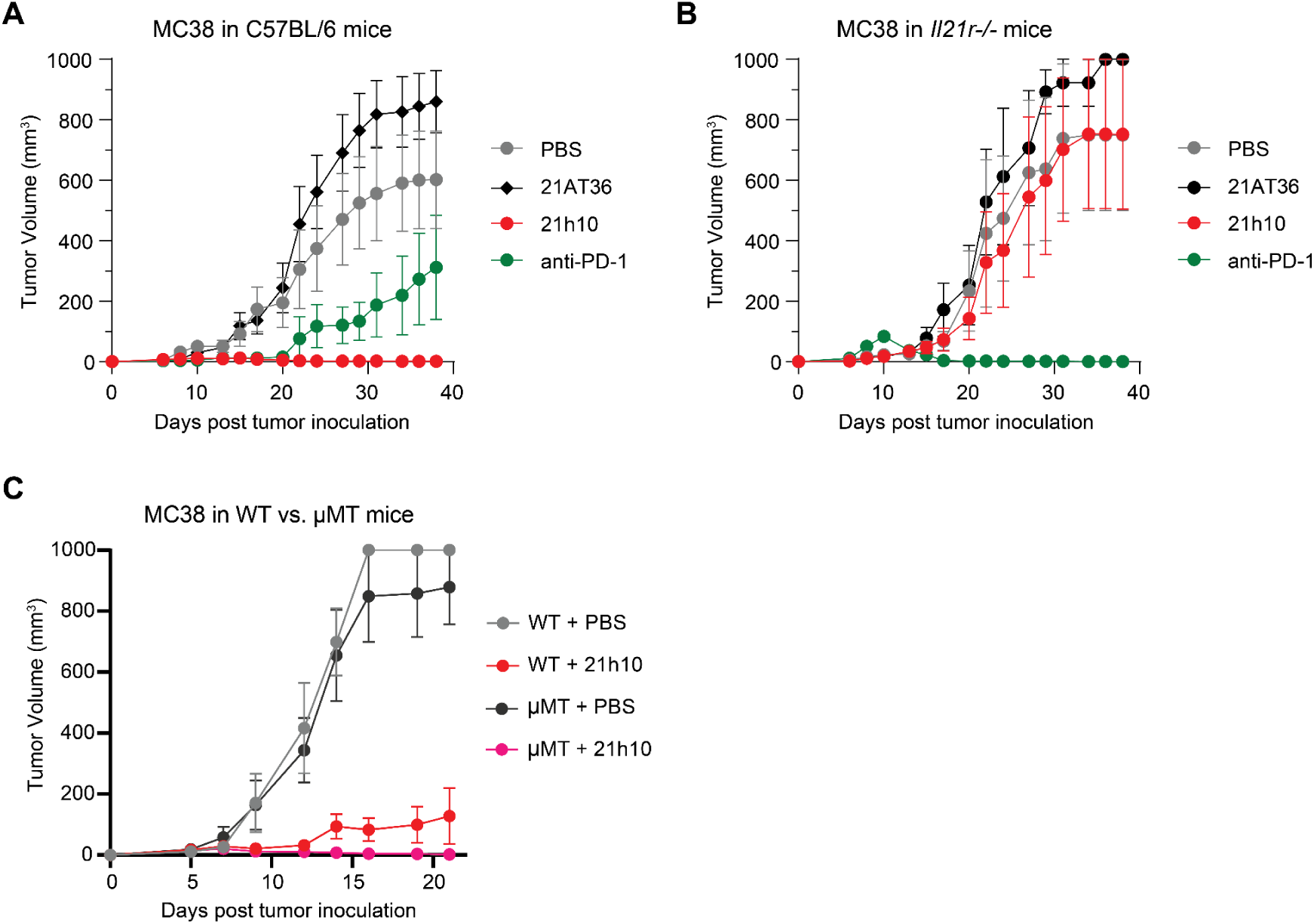
21h10 shows antitumor activity against MC38 adenocarcinoma in a T cell-dependent manner. (**A**) Antitumor activity of 21h10, 21AT36, and anti-PD-1 in murine MC38 in C57BL/6 mice, and (**B**) in *Il21r-/-* mice. (**C**) The antitumor activity of 21h10 in MC38 was compared in WT and μMT mice.

**Figure S7.**
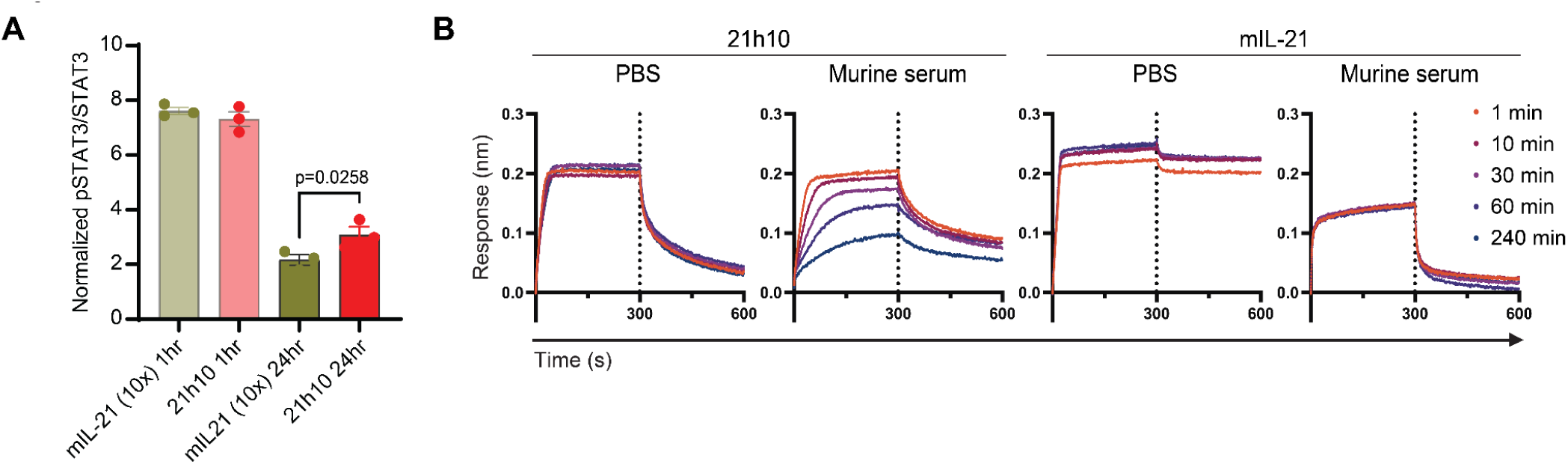
21h10 shows enhanced serum stability than mIL-21. (**A**) C57BL/6 mice were injected intravenously with a single dose of 21h10 or 10-fold molar excess of native mIL-21. Lymph nodes were harvested 1 or 24 hours later and analysed by immunoblot for phosphorylated STAT3. (**B**) BLI of mIL-21 and 21h10 to quantify the amount of protein binding to mIL-21R in PBS or murine serum for serum stability comparison. mIL-21 and 21h10 were incubated in PBS or murine serum for 1-240 minutes at 37°C. PBS incubation shows maximal binding possible with the given proteins. mIL-21 shows a rapid reduction of binding signal to almost half upon serum incubation compared to a gradual reduction in the case of 21h10. Maximal binding signals are shown in the groups with PBS incubation.

**Figure S8.**
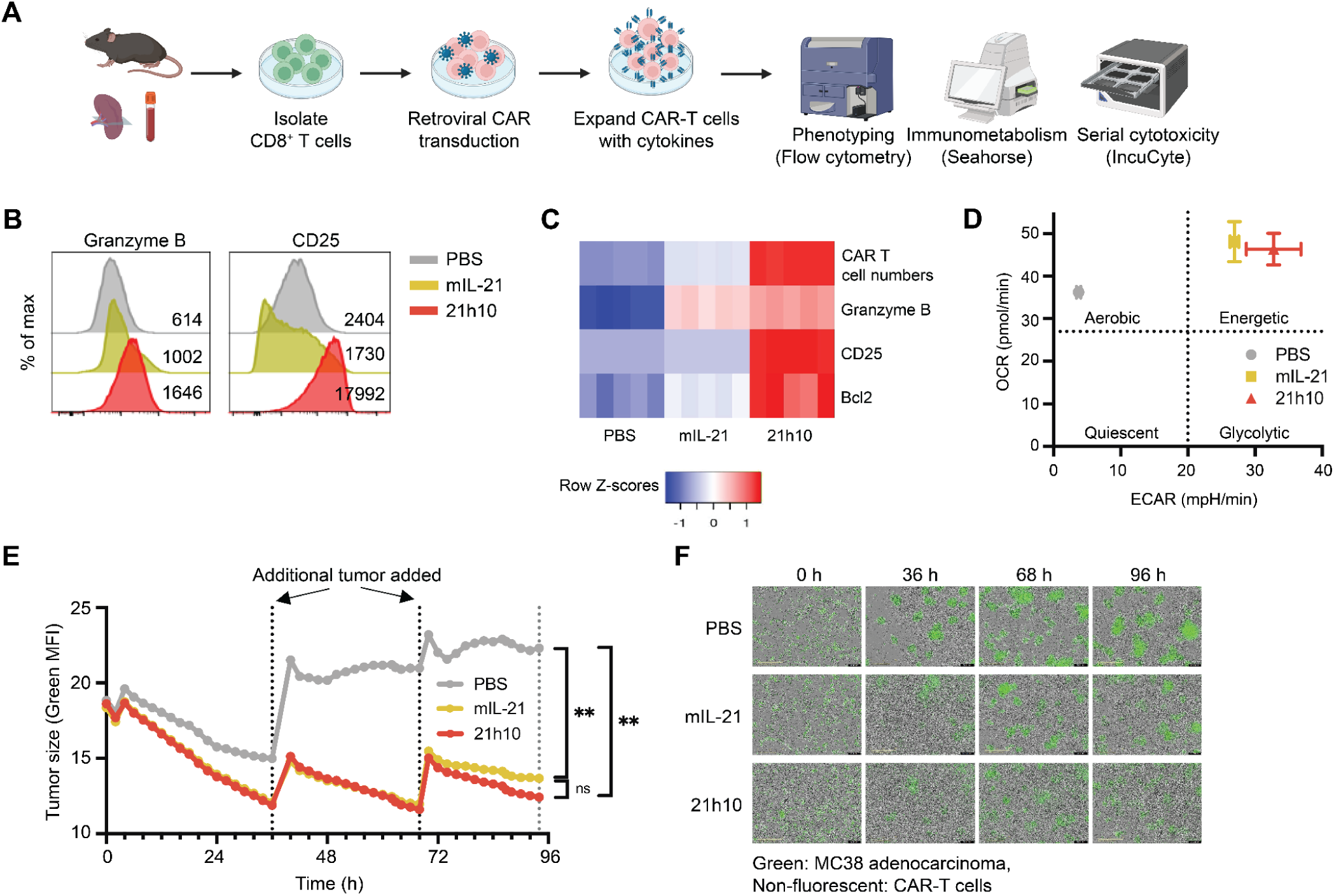
21h10 enhances cytotoxicity of CAR-T cells and increases their serial tumor-killing capability against MC38 adenocarcinoma. (**A**) CD8^+^ T cells were isolated and activated using anti-CD3/CD28, followed by transduction with a retroviral vector expressing CD19-targeting CAR to generate anti-CD19 CAR-T cells. (**B**) 21h10 increased the expression of granzyme B and CD25 in CAR-T cells, which increased with 21h10 treatment. Numbers in the histogram plots indicate the mean fluorescence intensity (MFI) of marker expression. (**C**) MFI of CAR expression, cell numbers, and MFI of granzyme B, CD25, and Bcl2 levels between PBS-, mIL-21-, and 21h10-treated CAR-T cells are presented as heat maps. (**D**) Metabolic profiles of PBS-, mIL-21-, and 21h10-treated CAR-T cells show potent induction of oxygen consumption rate (OCR) and extracellular acidification rate (ECAR) in Seahorse assay by 21h10. (**E**) Serial tumor killing of CD19^+^ GFP-MC38 adenocarcinoma by anti-CD19 CAR-T cells treated with PBS, mIL-21, or 21h10. (**F**) IncuCyte images of serial tumor killing assay in (**E**).

**Figure S9.**
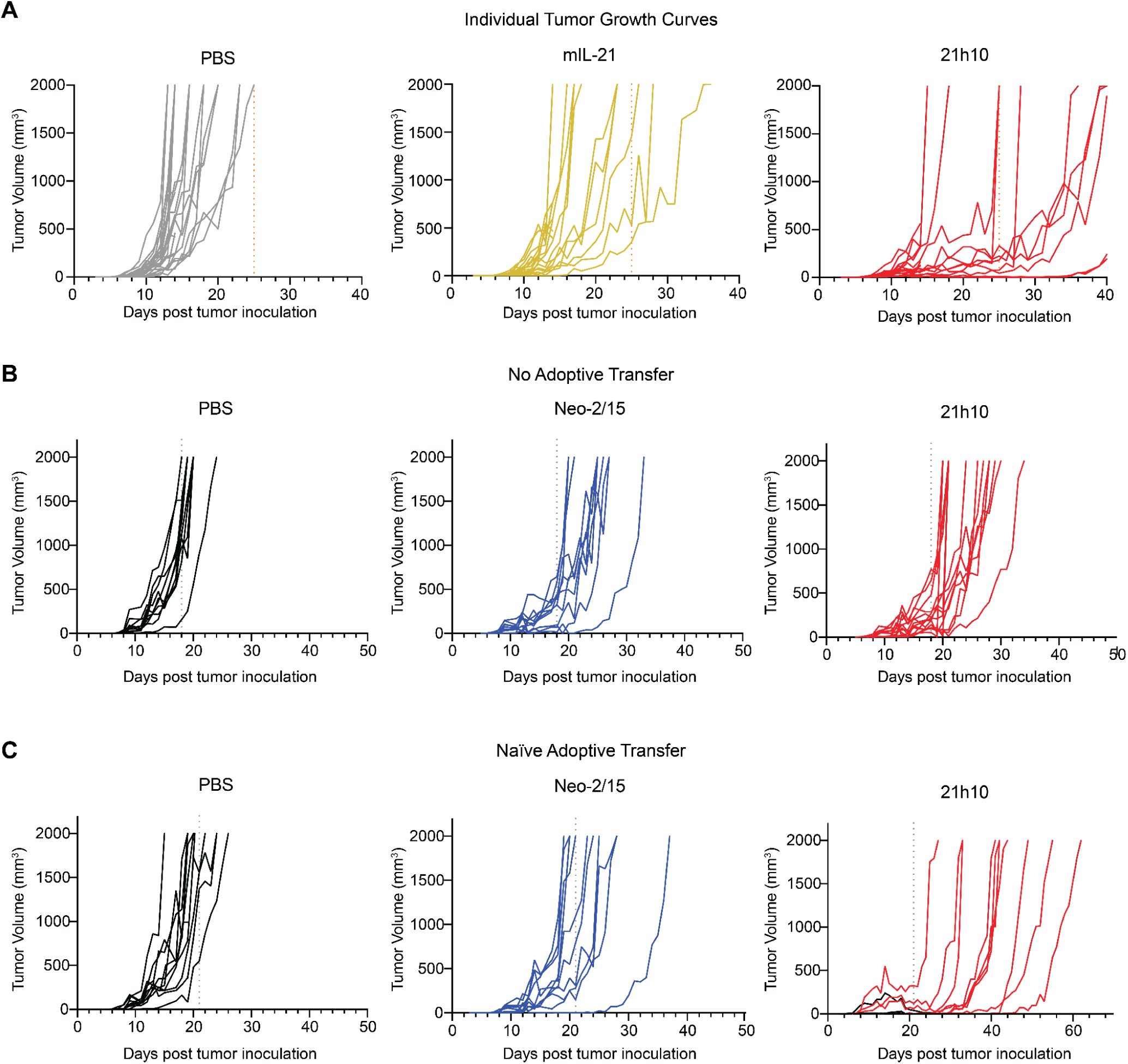
Individual tumor growth curves from. **Fig. 3**. (**A**) Individual tumor growth curves from Fig. 3A. (**B**) Individual tumor growth curves from Fig. 3B. (**C**) Individual tumor growth curves from Fig. 3C.

**Figure S10.**
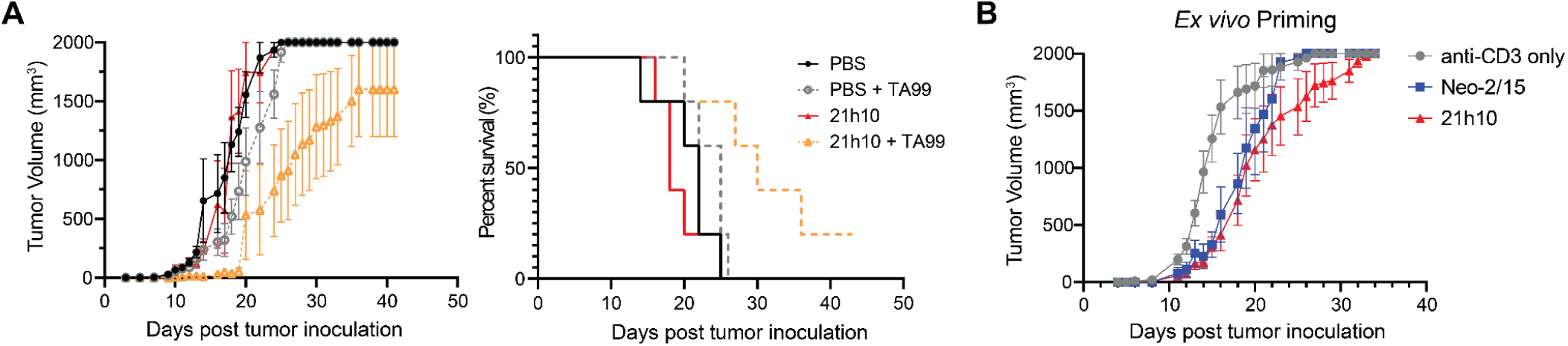
21h10 improves antibody therapy and *ex vivo*-primed T cell therapy. (**A**) Mice received B16F10 tumors and treatment with PBS or 21h10. Treatments were done alone or combined with TA99 (anti-TRP1 antibody). (**B**) TRP1 melanoma-specific T cells were primed *ex vivo* using anti-CD3 alone or in combination with Neo-2/15 or 21h10 for 2 days. Primed cells were transferred to mice 2 days after receiving B16F10 tumors.

**Figure S11.**
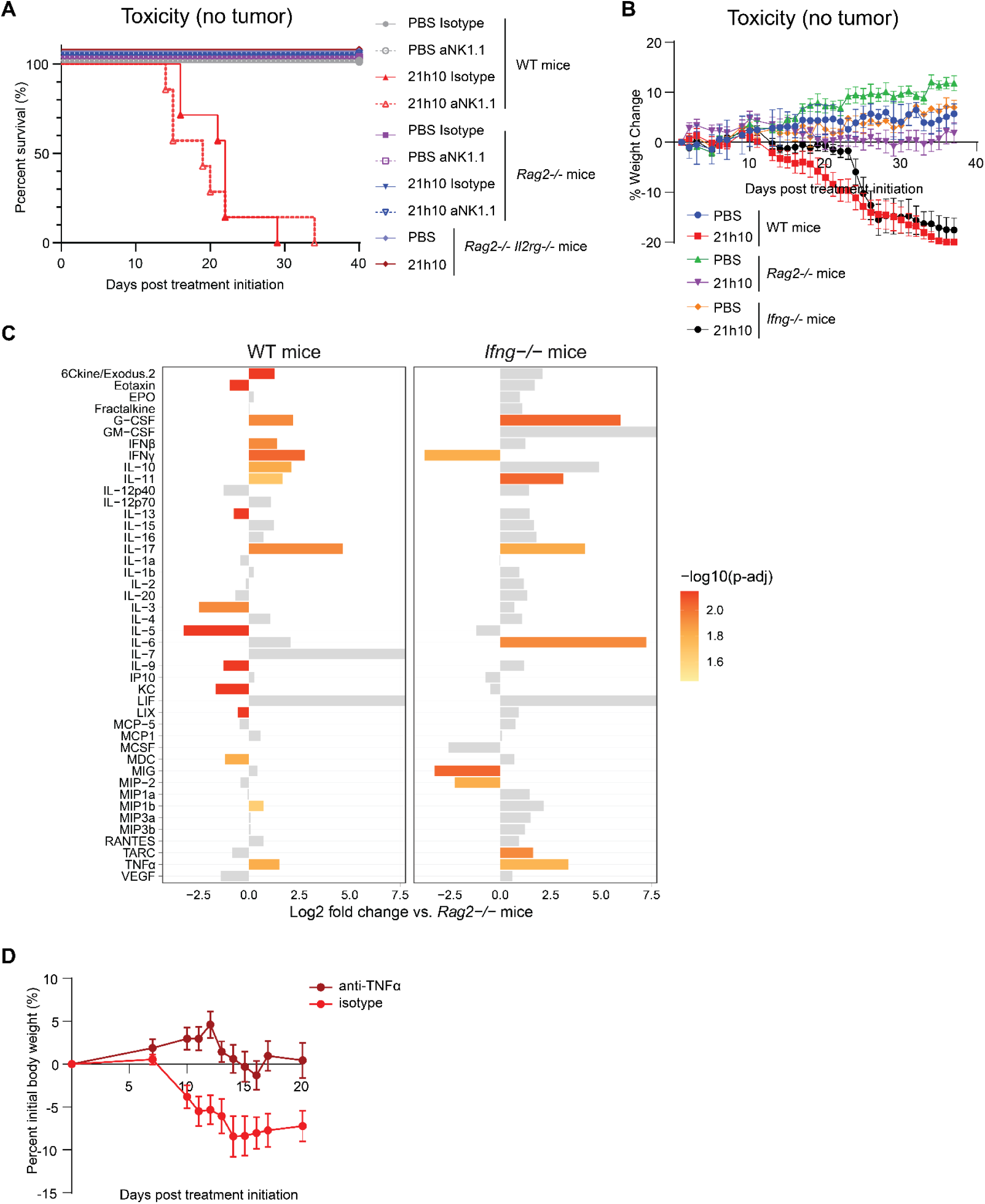
Toxicity profiling of 21h10 treatment. (**A**) Wild-type (WT), *Rag2-/-*, or *Rag2-/- Il2rg-/-* mice were treated with PBS or 21h10 daily and isotype or anti-NK1.1 depleting antibodies every three days. Mice were sacrificed when weight loss exceeded 20% of the initial starting weight. (**B**) WT, *Rag2-/-*, or *Ifng-/-* mice were treated with PBS or 21h10 daily. Mice were sacrificed when weight loss exceeded 20% of the initial starting weight. (**C**) Serum cytokine levels of various cytokines after 21h10 treatment in WT and *Ifng-/-* mice. Values plotted in comparison to *Rag2*-/- mice. Bars in gray are non-significant compared to *Rag2*-/- mice. (**D**) Body weight change plot in isotype- and anti-TNFα antibody- treated mice with 21h10 treatment.

**Figure S12.**
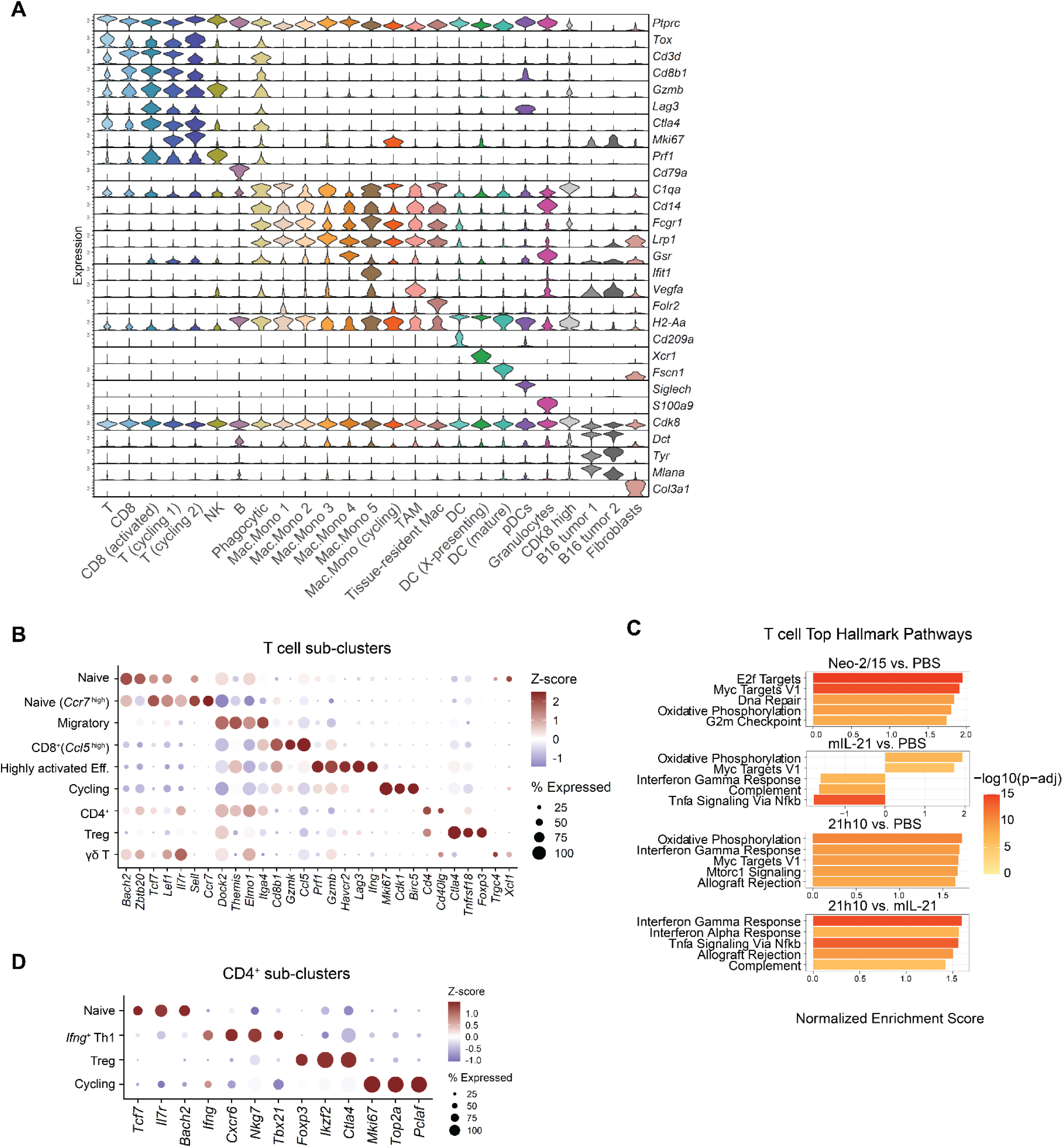
Cluster-defining genes for single-cell RNA-sequencing. (**A**) Violin plot of canonical gene expression for each cluster identified from Fig. 4A. (**B**, **D**) Dot plot of cluster-defining genes for T cell sub-clusters or CD4^+^ T cell sub-clusters from Fig. 4F or Fig. 4K, respectively. (**C**) Hallmark Gene Set Enrichment Analysis within T cells for given comparisons.

**Figure S13.**
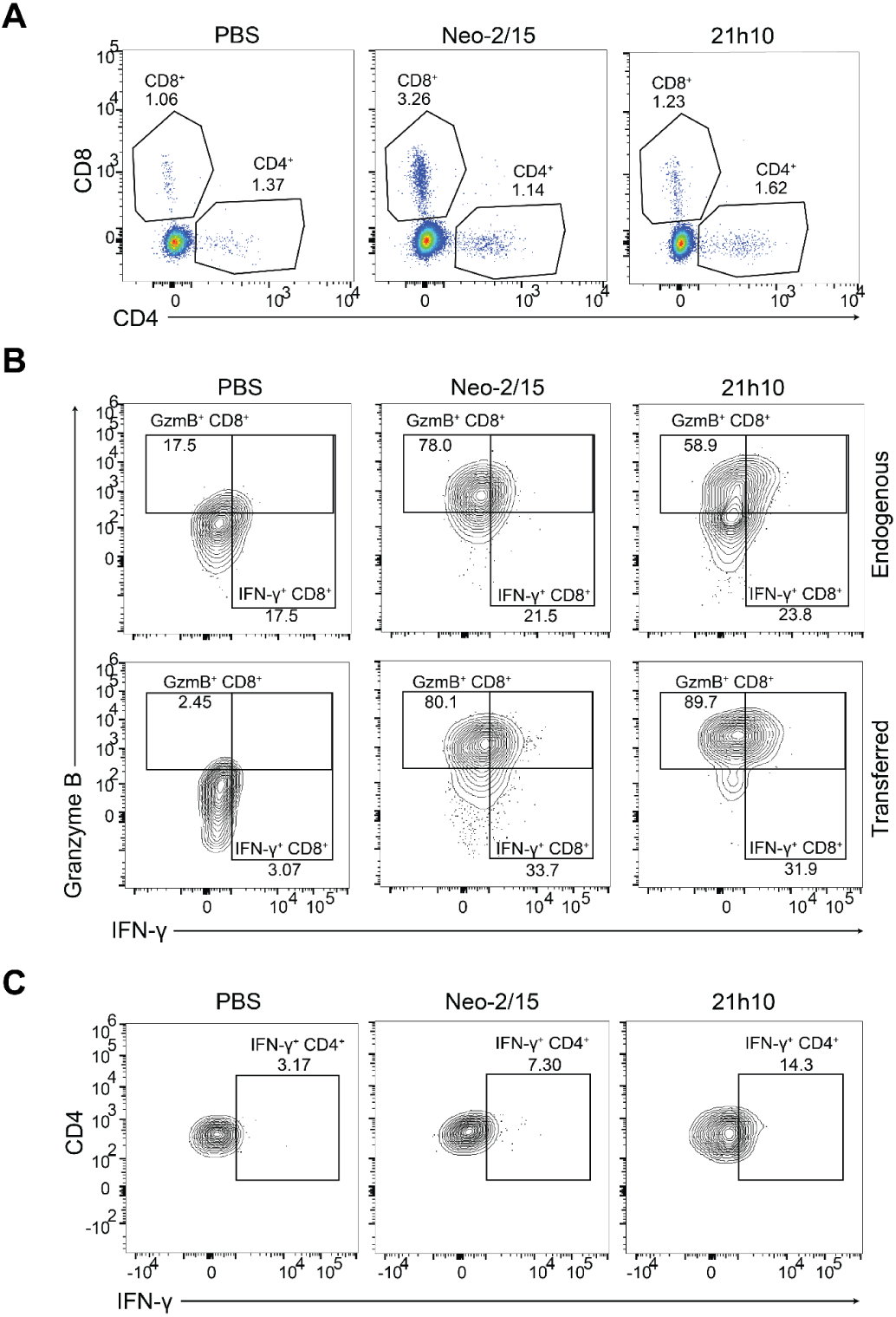
Representative flow cytometry plots. Representative flow cytometry plots from (**A**) Fig. 4D, (**B**) Fig. 4J, or (**C**) Fig. 4L.

**Figure S14.**
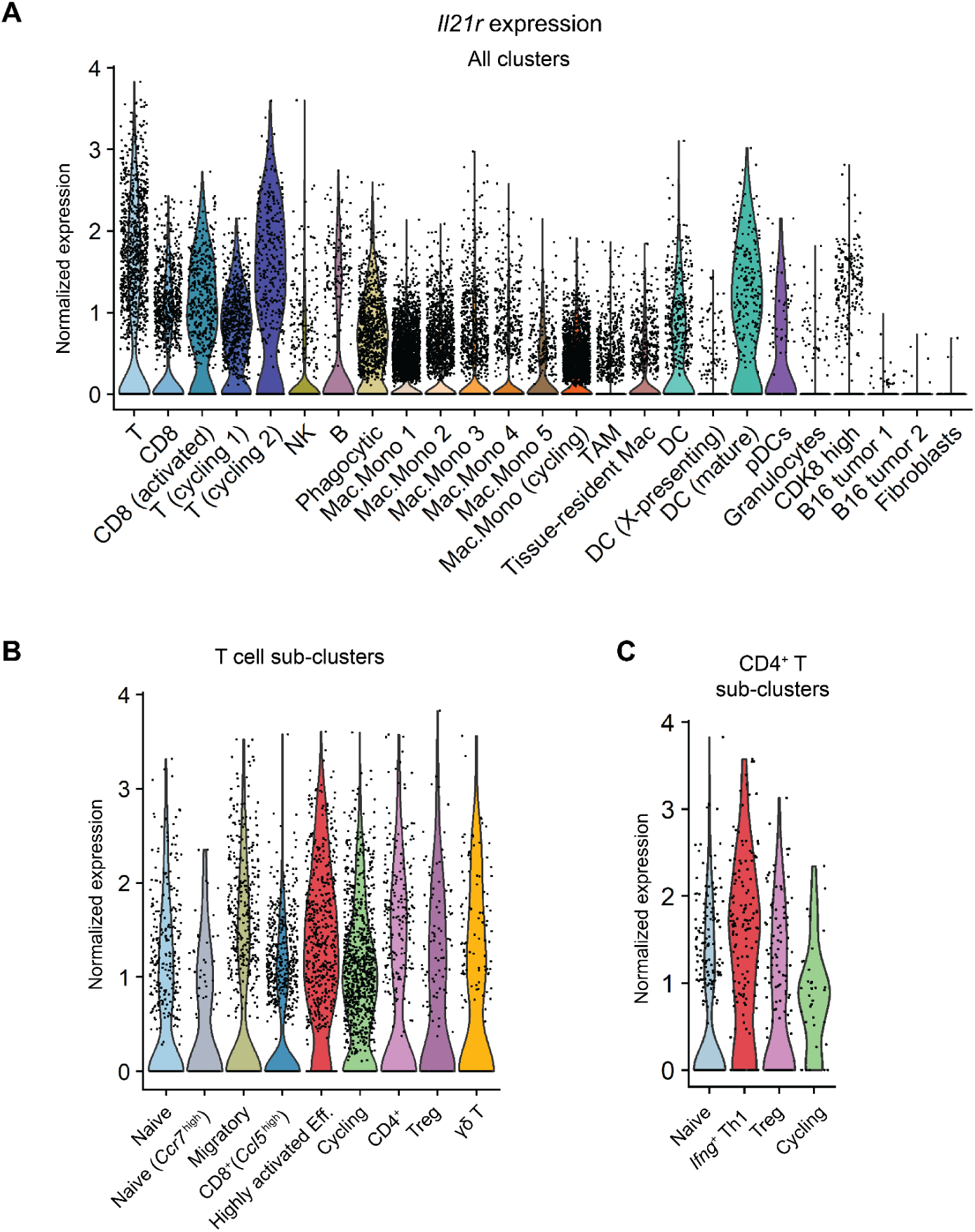
*Il21r* expression across clusters. (**A**) Violin plots of normalized *Il21r* expression for total clusters, (**B**) T cell sub-clusters, or (**C**) CD4^+^ T cell sub-clusters.

**Table S1.** K_D_ values of biolayer interferometry (BLI) assays.

| EC<sub>50</sub> (M) | Receptor | Kinetic fit values |  |  |  |  |
| --- | --- | --- | --- | --- | --- | --- |
| | | $K_D$ (Error) [M] | $k_{on}$ (Error) [M<sup>-1</sup>s<sup>-1</sup>] | $k_{off}$ (Error) [s<sup>-1</sup>] | Full X<sup>2</sup> | Full R<sup>2</sup> |
| 21h10 | hIL-21R | 4.82E-10 (7.63E-11) | 2.20E+05 (3.08E+03) | 1.06E-04 (1.67E-05) | 2.051 | 0.9812 |
|  | hIL-21R/hγ<sub>c</sub> | 8.67E-08 (1.90E-09) | 7.45E+04 (1.53E+03) | 6.46E-03 (5.06E-05) | 0.2295 | 0.9813 |
|  | mIL-21R | 7.87E-09 (1.25E-10) | 3.92E+05 (5.59E+03) | 3.09E-03 (2.13E-05) | 0.9389 | 0.9750 |
|  | mIL-21R/mγ<sub>c</sub> | 7.28E-08 (4.30E-09) | 1.14E+05 (6.35E+03) | 8.29E-03 (1.62E-04) | 0.1802 | 0.9232 |
| 21AT36 | hIL-21R | 9.19E-10 (1.06E-10) | 2.05E+05 (3.78E+03) | 1.88E-04 (2.14E-05) | 1.1895 | 0.9763 |
|  | mIL-21R | 1.05E-08 (2.13E-10) | 3.08E+05 (5.66E+03) | 3.24E-03 (2.83E-05) | 0.4466 | 0.9747 |

**Table S2.**
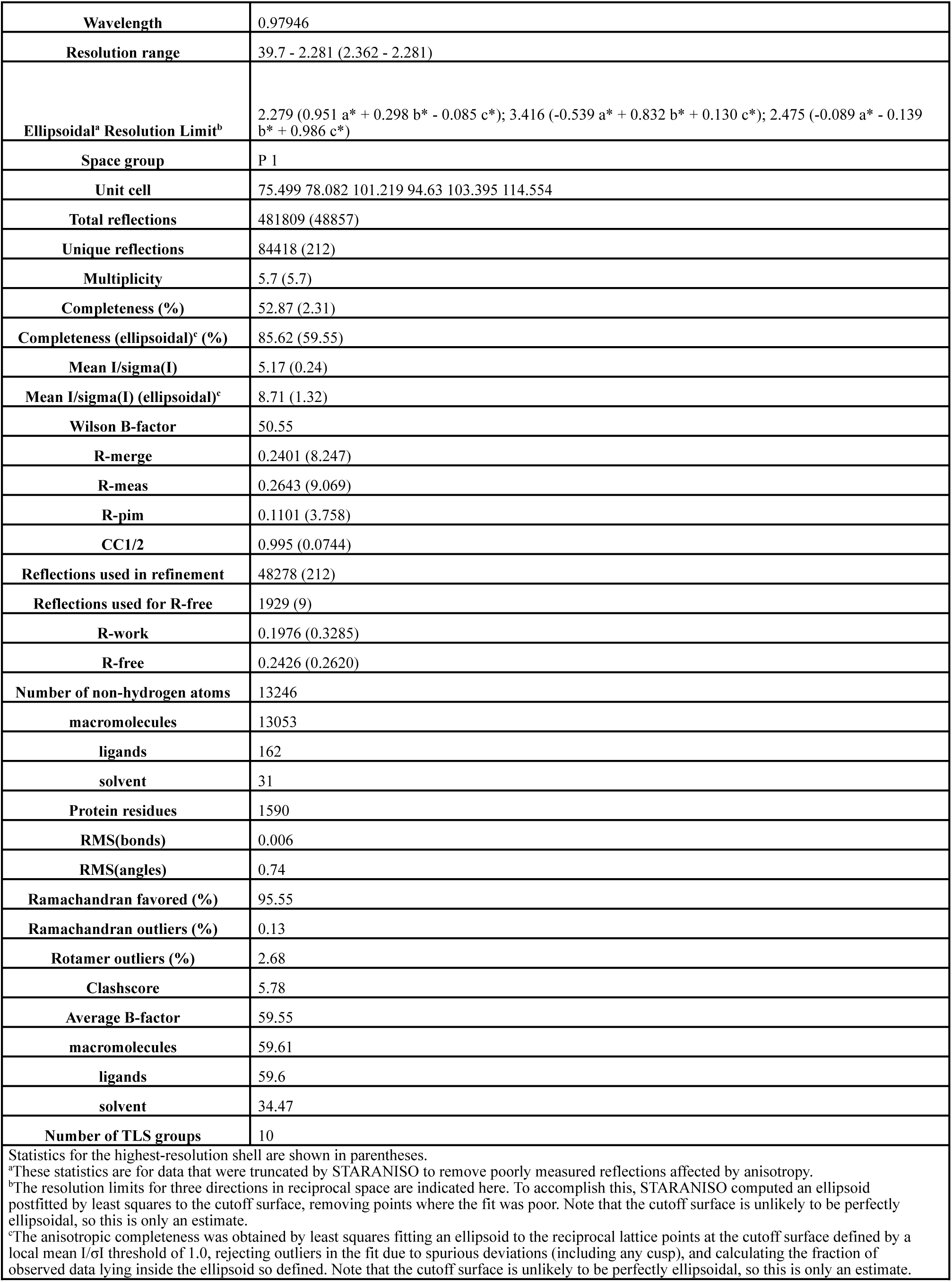
Crystallography data collection and refinement statistics.

**Table S3.** Key contacts in the 21h10 receptor complex.

| Interface overview |  |  |  | Site IIa<br>21h10-h<sub>γc</sub> interface |  |  |  |
| --- | --- | --- | --- | --- | --- | --- | --- |
| Interaction | Structure 1 | Structure 2 | Interface [Å<sup>2</sup>] | Hydrogen bonds | h<sub>γc</sub> | Distance [Å] | 21h10 |
| Site I | 21h10 | hIL-21R | 870.7 | 1 | C:GLN149[OE1] | 3.08 | A:SER92[OG] |
| Site IIa | 21h10 | h<sub>γc</sub> | 423.1 | 2 | C:GLN149[OE1] | 3.10 | A:GLN95[NE2] |
| Site IIb | hIL-21R | h<sub>γc</sub> | 693.8 |  |  |  |  |

| Site I<br>21h10-hIL-21R interface |  |  |  | Site IIb<br>hIL-21R-h<sub>γc</sub> interface |  |  |  |
| --- | --- | --- | --- | --- | --- | --- | --- |
| Hydrogen bonds | hIL-21R | Distance [Å] | 21h10 | Hydrogen bonds | h<sub>γc</sub> | Distance [Å] | hIL-21R |
| 1 | B:ASP72[O] | 2.78 | A:ARG59[NH2] | 1 | C:ARG205[NH1] | 2.84 | B:GLN136[OE1] |
| 2 | B:ASP72[OD2] | 2.91 | A:ARG63[NH1] | 2 | C:SER212[OG] | 3.25 | B:SER165[O] |
| 3 | B:ASP72[OD2] | 3.15 | A:ARG46[NH1] | 3 | C:SER168[O] | 2.99 | B:LYS155[NZ] |
| 4 | B:MET130[SD] | 3.25 | A:LYS39[NZ] | 4 | C:GLN169[OE1] | 2.48 | B:LEU156[N] |
| 5 | B:GLY188[O] | 3.28 | A:ARG65[NH2] |  |  |  |  |

| Site I<br>21h10-hIL-21R interface |  |  |  |
| --- | --- | --- | --- |
| Salt bridges | hIL-21R | Distance [Å] | 21h10 |
| 1 | B:LYS134[NZ] | 3.77 | A:ASP69[OD1] |
| 2 | B:LYS134[NZ] | 3.39 | A:ASP69[OD2] |
| 3 | B:ASP72[OD2] | 2.91 | A:ARG63[NH1] |
| 4 | B:ASP72[OD2] | 3.15 | A:ARG46[NH1] |
| 5 | B:ASP73[OD1] | 3.10 | A:ARG46[NH1] |
| 6 | B:ASP73[OD2] | 3.73 | A:ARG46[NH2] |
| 7 | B:ASP73[OD2] | 3.36 | A:ARG42[NH2] |

**Table S4.** Mean EC_50_ values from cross-reactivity cell signaling assays.

| EC<sub>50</sub> (M) | Human CD8<sup>+</sup> T cells |  | Murine CD8<sup>+</sup> T cells |  |
| --- | --- | --- | --- | --- |
|  | STAT1 | STAT3 | STAT1 | STAT3 |
| **hIL-21** | 4.577E-11 | 4.062E-11 | 1.179E-08 | 5.563E-10 |
| **mIL-21** | 3.563E-11 | 2.964E-11 | 6.227E-11 | 1.645E-11 |
| **21h10** | 9.990E-12 | 1.030E-11 | 1.592E-11 | 5.684E-12 |

**Table S5.** Mean EC_50_ values of cell signaling assays.

| EC<sub>50</sub> (M) | Human CD8<sup>+</sup> T cells |  | Murine CD8<sup>+</sup> T cells |  |
| --- | --- | --- | --- | --- |
|  | STAT1 | STAT3 | STAT1 | STAT3 |
| **hIL-21** | 5.593E-11 | 3.946E-11 | N/M | N/M |
| **mIL-21** | N/M | N/M | 5.073E-11 | 4.153E-11 |
| **21h10** | 2.347E-11 | 1.475E-11 | 4.560E-11 | 2.516E-11 |

| EC<sub>50</sub> (M) | Human CD4<sup>+</sup> T cells |  | Murine CD4<sup>+</sup> T cells |  |
| --- | --- | --- | --- | --- |
|  | STAT1 | STAT3 | STAT1 | STAT3 |
| **hIL-21** | 1.447E-11 | 8.991E-12 | N/M | N/M |
| **mIL-21** | N/M | N/M | 4.523E-11 | 3.675E-11 |
| **21h10** | 6.624E-12 | 3.470E-12 | 7.015E-11 | 3.917E-11 |

| EC<sub>50</sub> (M) | Human B cells |  | Murine B cells |  |
| --- | --- | --- | --- | --- |
|  | STAT1 | STAT3 | STAT1 | STAT3 |
| **hIL-21** | 2.348E-11 | 9.064E-12 | N/M | N/M |
| **mIL-21** | N/M | N/M | 4.198E-11 | 6.517E-12 |
| **21h10** | 1.233E-11 | 2.078E-12 | 9.095E-12 | 2.865E-12 |

\*N/M: not measured

**Table S6.** Clinical, demographic, and molecular characteristics of patient samples analyzed in the ex vivo PDOTS experiment.

| Sample # | Tumor Type | Age | Gender | Mutational Status | Tumor mutation burden (absolute count) | Stage | R/NR |
| --- | --- | --- | --- | --- | --- | --- | --- |
| 10368 | cutaneous melanoma | 91 | F | N/A | N/A | N/A | NR |
| 10370 | cutaneous melanoma | 51 | M | BRAFV600E, TERT, EGFR | none | IV | NR |
| 10373 | Merkel cell carcinoma | 83 | F | N/A | N/A | N/A | NR |
| 10387 | cutaneous melanoma | 35 | F | CDKN2A, TERT, BRAF | low | III | NR |
| 10388 | cutaneous melanoma | 99 | M | none | none | III | Tx naïve |
| 10406 | cutaneous melanoma | 49 | M | N/A | N/A | N/A | N/A |
| 10407 | cutaneous melanoma | 86 | M | none | N/A | III | NR |
| 10443 | cutaneous melanoma | 57 | M | PTEN, CDKN2A, TERT and KEAP1, BRAF-WT | low | IV | NR |
| 10488 | cutaneous melanoma | 75 | M | TERT and BRAF | low | IV | Tx naïve |
| 10450 | cutaneous melanoma | 69 | M | NRAS, FBXW7 | low | III | NR |
| 10451 | cutaneous melanoma | 41 | M | PTEN, CDKN2A, TERT and BRAF | low | III | NR |
| 10452 | cutaneous melanoma | 69 | M | DAXX, BRAF and KRAS | low | III | NR |
| 10456 | Acral melanoma | 80 | F | CCND2, CDKN2A and NF1 | low | III | NR |

